# Evolutionary constraint and innovation across hundreds of placental mammals

**DOI:** 10.1101/2023.03.09.531574

**Authors:** Matthew J. Christmas, Irene M. Kaplow, Diane P. Genereux, Michael X. Dong, Graham M. Hughes, Xue Li, Patrick F. Sullivan, Allyson G. Hindle, Gregory Andrews, Joel C. Armstrong, Matteo Bianchi, Ana M. Breit, Mark Diekhans, Cornelia Fanter, Nicole M. Foley, Daniel B. Goodman, Linda Goodman, Kathleen C. Keough, Bogdan Kirilenko, Amanda Kowalczyk, Colleen Lawless, Abigail L. Lind, Jennifer R. S. Meadows, Lucas R. Moreira, Ruby W. Redlich, Louise Ryan, Ross Swofford, Alejandro Valenzuela, Franziska Wagner, Ola Wallerman, Ashley R. Brown, Joana Damas, Kaili Fan, John Gatesy, Jenna Grimshaw, Jeremy Johnson, Sergey V. Kozyrev, Alyssa J. Lawler, Voichita D. Marinescu, Kathleen M. Morrill, Austin Osmanski, Nicole S. Paulat, BaDoi N. Phan, Steven K. Reilly, Daniel E. Schäffer, Cynthia Steiner, Megan A. Supple, Aryn P. Wilder, Morgan E. Wirthlin, James R. Xue, Zoonomia Consortium, Bruce W. Birren, Steven Gazal, Robert M. Hubley, Klaus-Peter Koepfli, Tomas Marques-Bonet, Wynn K. Meyer, Martin Nweeia, Pardis C. Sabeti, Beth Shapiro, Arian F. A. Smit, Mark Springer, Emma Teeling, Zhiping Weng, Michael Hiller, Danielle L. Levesque, Harris A. Lewin, William J. Murphy, Arcadi Navarro, Benedict Paten, Katherine S. Pollard, David A. Ray, Irina Ruf, Oliver A. Ryder, Andreas R. Pfenning, Kerstin Lindblad-Toh, Elinor K. Karlsson

**Affiliations:** Department of Medical Biochemistry and Microbiology, Science for Life Laboratory, Uppsala University; Uppsala, 751 32, Sweden; Department of Computational Biology, School of Computer Science, Carnegie Mellon University; Pittsburgh, PA 15213, USA; Neuroscience Institute, Carnegie Mellon University; Pittsburgh, PA 15213, USA; Broad Institute of MIT and Harvard; Cambridge, MA 02139, USA; School of Biology and Environmental Science, University College Dublin; Belfield, Dublin 4, Ireland; Morningside Graduate School of Biomedical Sciences, UMass Chan Medical School; Worcester, MA 01605, USA; Program in Bioinformatics and Integrative Biology, UMass Chan Medical School; Worcester, MA 01605, USA; Department of Genetics, University of North Carolina Medical School; Chapel Hill, NC 27599, USA; Department of Medical Epidemiology and Biostatistics, Karolinska Institutet; Stockholm, Sweden; School of Life Sciences, University of Nevada Las Vegas; Las Vegas, NV 89154, USA; Genomics Institute, University of California Santa Cruz; Santa Cruz, CA 95064, USA; School of Biology and Ecology, University of Maine; Orono, ME 04469, USA; Veterinary Integrative Biosciences, Texas A&M University; College Station, TX 77843, USA; Fauna Bio Incorporated; Emeryville, CA 94608, USA; Department of Epidemiology & Biostatistics, University of California San Francisco; San Francisco, CA 94158, USA; Gladstone Institutes; San Francisco, CA 94158, USA; Faculty of Biosciences, Goethe-University; 60438 Frankfurt, Germany; LOEWE Centre for Translational Biodiversity Genomics; 60325 Frankfurt, Germany; Senckenberg Research Institute; 60325 Frankfurt, Germany; Department of Biological Sciences, Mellon College of Science, Carnegie Mellon University; Pittsburgh, PA 15213, USA; Department of Experimental and Health Sciences, Institute of Evolutionary Biology (UPF-CSIC), Universitat Pompeu Fabra; Barcelona, 08003, Spain; Museum of Zoology, Senckenberg Natural History Collections Dresden; 01109 Dresden, Germany; The Genome Center, University of California Davis; Davis, CA 95616, USA; Division of Vertebrate Zoology, American Museum of Natural History; New York, NY 10024, USA; Department of Biological Sciences, Texas Tech University; Lubbock, TX 79409, USA; Medical Scientist Training Program, University of Pittsburgh School of Medicine; Pittsburgh, PA 15261, USA; Department of Genetics, Yale School of Medicine; New Haven, CT 06510, USA; Conservation Genetics, San Diego Zoo Wildlife Alliance; Escondido, CA 92027, USA; Department of Ecology and Evolutionary Biology, University of California Santa Cruz; Santa Cruz, CA 95064, USA; Allen Institute for Brain Science; Seattle, WA 98109, USA; Department of Organismic and Evolutionary Biology, Harvard University; Cambridge, MA 02138, USA; Keck School of Medicine, University of Southern California; Los Angeles, CA 90033, USA; Institute for Systems Biology; Seattle, WA 98109, USA; Center for Species Survival, Smithsonian’s National Zoo and Conservation Biology Institute; Washington, DC 20008, USA; Computer Technologies Laboratory, ITMO University; St. Petersburg 197101, Russia; Smithsonian-Mason School of Conservation, George Mason University; Front Royal, VA 22630, USA; Catalan Institution of Research and Advanced Studies (ICREA); Barcelona, 08010, Spain; CNAG-CRG, Centre for Genomic Regulation, Barcelona Institute of Science and Technology (BIST); Barcelona, 08036, Spain; Department of Medicine and LIfe Sciences, Institute of Evolutionary Biology (UPF-CSIC), Universitat Pompeu Fabra; Barcelona, 08003, Spain; Institut Catalàde Paleontologia Miquel Crusafont, Universitat Autònoma de Barcelona; 08193, Cerdanyola del Vallès, Barcelona, Spain; Department of Biological Sciences, Lehigh University; Bethlehem, PA 18015, USA; Department of Comprehensive Care, School of Dental Medicine, Case Western Reserve University; Cleveland, OH 44106, USA; Department of Vertebrate Zoology, Canadian Museum of Nature; Ottawa, Ontario K2P 2R1, Canada; Department of Vertebrate Zoology, Smithsonian Institution; Washington, DC 20002, USA; Narwhal Genome Initiative, Department of Restorative Dentistry and Biomaterials Sciences, Harvard School of Dental Medicine; Boston, MA 02115, USA; Howard Hughes Medical Institute; Chevy Chase, MD, USA; Howard Hughes Medical Institute, University of California Santa Cruz; Santa Cruz, CA 95064, USA; Department of Evolution, Ecology and Organismal Biology, University of California Riverside; Riverside, CA 92521, USA; Department of Evolution and Ecology, University of California Davis; Davis, CA 95616, USA; John Muir Institute for the Environment, University of California Davis; Davis, CA 95616, USA; BarcelonaBeta Brain Research Center, Pasqual Maragall Foundation; Barcelona, 08005, Spain; CRG, Centre for Genomic Regulation, Barcelona Institute of Science and Technology (BIST); Barcelona, 08003, Spain; Chan Zuckerberg Biohub; San Francisco, CA 94158, USA; Division of Messel Research and Mammalogy, Senckenberg Research Institute and Natural History Museum Frankfurt; 60325 Frankfurt am Main, Germany; Department of Evolution, Behavior and Ecology, School of Biological Sciences, University of California San Diego; La Jolla, CA 92039, USA; Program in Molecular Medicine, UMass Chan Medical School; Worcester, MA 01605, USA

## Abstract

Evolutionary constraint and acceleration are powerful, cell-type agnostic measures of functional importance. Previous studies in mammals were limited by species number and reliance on human-referenced alignments. We explore the evolution of placental mammals, including humans, through reference-free whole-genome alignment of 240 species and protein-coding alignments for 428 species. We estimate 10.7% of the human genome is evolutionarily constrained. We resolve constraint to single nucleotides, pinpointing functional positions, and refine and expand by over seven-fold the catalog of ultraconserved elements. Overall, 48.5% of constrained bases are as yet unannotated, suggesting yet-to-be-discovered functional importance. Using species-level phenotypes and an updated phylogeny, we associate coding and regulatory variation with olfaction and hibernation. Focusing on biodiversity conservation, we identify genomic metrics that predict species at risk of extinction.

## Main text

Placental mammals, the evolutionary lineage that includes humans, are exceptionally diverse. With more than 5,000 extant species, from the 2 gram bumblebee bat to the 150,000 kilogram blue whale, mammals have adapted to almost every habitat on earth^1, 2^. How rapidly placental mammals diversified and whether this preceded or followed the major extinction event at the end of the Cretaceous period(**Fig. 1A**), have been difficult to determine due to the dearth of fossil and molecular data^3, 4^.

**Figure 1:**
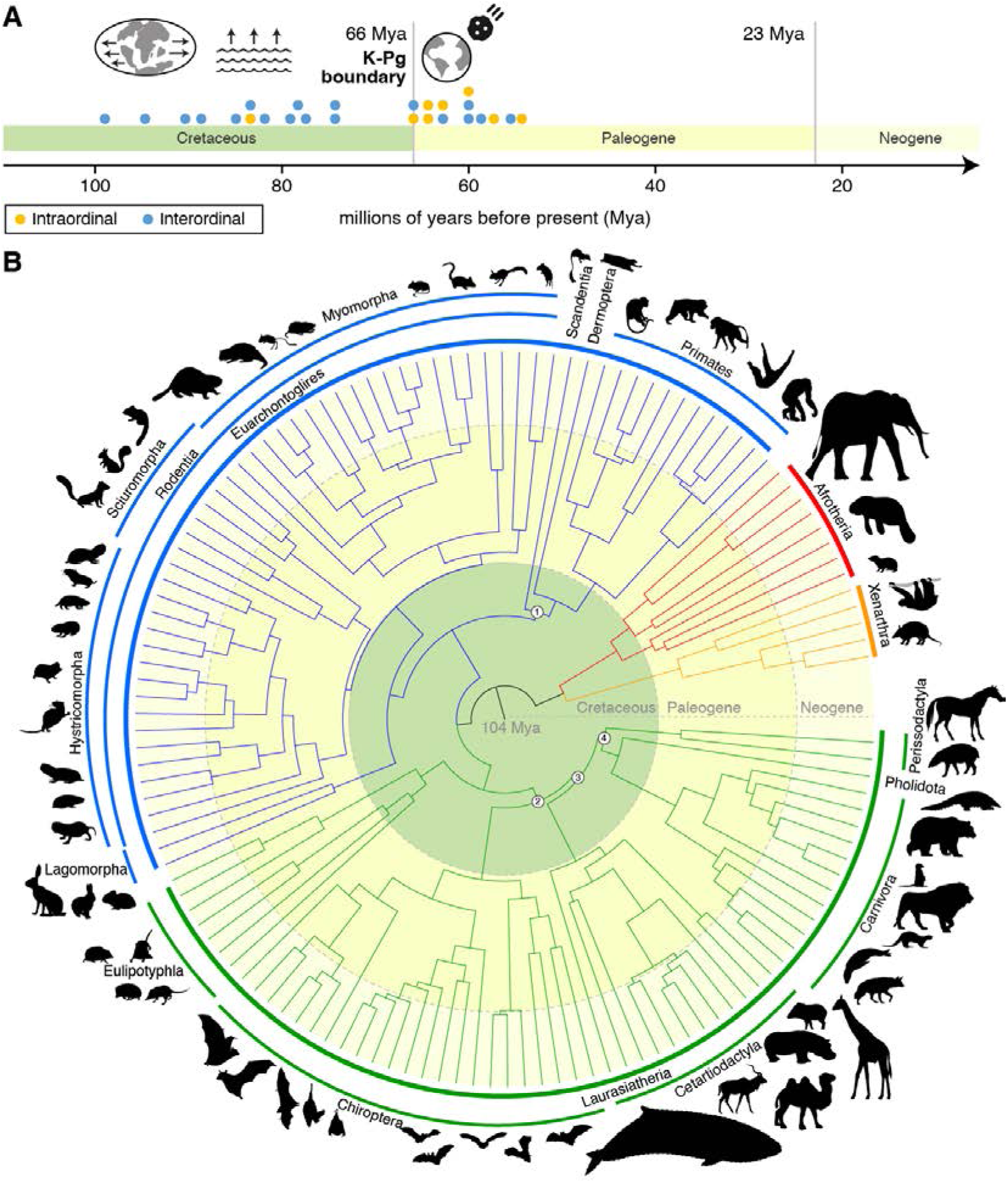
New placental mammal phylogeny supports Long Fuse model of diversification. **(A)** The majority of interordinal diversification (blue circles) occurred in the Cretaceous, coincident with continental fragmentation and sea level changes. A second pulse of intra-ordinal diversification (yellow circles) occured after the mass extinction event at the K-Pg boundary. **(B)** A phylogeny based on divergence times estimated using ∼460kb of near-neutrally evolving sequence for 240 species resolves recalcitrant relationships in the placental mammal phylogeny (black numbers in white circles) including(1) Euarchonta (primates, colugos, and treeshrews);(2) Scrotifera (bats, cetartiodactyls, perissodactyls, carnivorans, and pangolins);(3) Feruungulata (all scrotiferans excluding bats); and(4) Ferae (carnivorans and pangolins) + Perissodactyla.

By comparing genomes, we can find positions whose unusual patterns of variation suggest functional importance^5^. Lack of variation (conservation) suggests purifying selection. Higher than expected variation in specific lineages (acceleration) suggests positive selection.

Exceptionally high conservation points to evolutionary constraint. Genomes also evolve through larger structural changes, including introgression and translocation of transposable elements^6^, deletions, and duplications^7^.

Associating evolutionary signals with species-level phenotypes can identify genetic variants underlying specific adaptations^8^, such as diet type^9^, echolocation^10^, and subterranean habitation^11^. Coding regions tend to change more slowly than regulatory regions, which frequently arise and disappear within placental lineages, and at which nucleotide-level conservation is sometimes weak ^12^. The sensitivity and specificity of methods for detecting evolutionary signals and associating them with phenotypes depends on the number and diversity of species analyzed^13, 14^.

Zoonomia is the largest mammalian comparative genomics resource produced to date, with whole genomes aligned for 240 species (2.3-fold more families and ∼4-fold more species than the mammals included in the earlier 100 Vertebrates alignment^15^) and protein-coding sequences aligned for 428 species. Our first publication described how species were selected, sequenced, and aligned using a reference-free approach that captures sequences missing from the human genome^16^. In a companion paper, we explore how evolutionary constraint is a powerful tool for mapping functional variants involved in human diseases^17^. Here, we address long-standing questions about the evolution of placental mammals, annotate regions of exceptional constraint and acceleration in the human genome, and investigate the origins of intriguing mammalian traits.

### New resources for comparative genomics

Zoonomia includes two large alignments: the 241-way reference-free Cactus alignment, with 240 species (domestic dog has two representatives)^16^, and the 428-way human-referenced TOGA (Tool to infer Orthologs from Genome Alignments) alignment of protein-coding sequences (19,464 genes)^18^. We measured sequence conservation using phyloP for versions of the Cactus alignment projected onto the human, chimpanzee, mouse, dog, and little brown bat reference genomes(**Table S1**)^5^. For the primate lineage (43 species), we also measured conservation using phastCons, which models multibase elements and achieves more power with fewer species^19^. In **Table S2**, we list all species included with their common and binomial names, associated analyses, and phenotypes.

### Phylogeny supports the “Long Fuse” Model

We used the Cactus alignment to infer a new phylogeny of placental mammals from near-neutrally evolving variable sites(N=466,232)(**Fig. 1B**)(detailed in^20^). Divergence time analyses supported the “Long Fuse” model of mammalian diversification, where interordinal diversification took place in the Cretaceous and the majority of interordinal diversification took place after the Cretaceous-Paleogene (K-Pg) mass extinction event(Fig. 1A)^21^. In contrast, our analyses did not support the fossil record-derived “Explosive” model, which places all inter- and intraordinal diversification after the K-Pg event, and alternative scenarios posited by numerous analyses of evolutionarily constrained sequences^22, 23^.

The timetree and phylogenomic analysis, displaying a relative lack of phylogenomic discordance at interordinal nodes, suggests interordinal diversification occurred via allopatry during a period of continental fragmentation and sea level change prior to the K-Pg boundary (**Fig. 1A**)^24, 25^. Conversely, some post-K-Pg intraordinal diversification events show high phylogenomic discordance, consistent with a history of incomplete lineage sorting and gene flow. Notably, we highlight a pulse of predominantly intraordinal diversification immediately following the K-Pg boundary (**Fig. 1A**), a molecular result consistent with the fossil record^26^.

### Reconstructing the mammalian ancestor

We reconstructed a karyotype for the ancestor of all mammals (placental, marsupial, and monotreme) using high-contiguity genome assemblies for 32 species, representing all but four orders (detailed in ^27^). Our sequence-based approach assigns 87% of the human genome to 20 ancestral chromosomes. We detected 365 rearrangements (145 new) and dated each on the mammalian phylogeny. On average ∼2 breakpoints occur every million years (My), but this varies widely. There are forty-fold more on the branch to the therian ancestor (3.9/My) than between the placental and boreoeutherian ancestor (0.1/My). Nearly half the human genome is in synteny blocks >1Mb and shared across all mammals, with the longest (22.1 Mb) encompassing the *HOXD* gene cluster and other genes involved in embryonic morphogenesis(**Table S3**).

### Transposable element activity

We cataloged transposable elements (TEs) in the genome assemblies for 243 species, producing by far the largest resource for investigating how these powerful agents of genomic change vary in placental mammals (detailed in ^28^)(**Fig. S1**). TEs are repetitive, mobile DNA sequences that tend to insert as units of 100 to 10,000bp and accumulate (>1 million copies per genome). We found that TEs comprise 49.0%±7.5% of each genome on average with little variability across species, consistent with a model where TE accumulation is counterbalanced by DNA loss^29^. Recent accumulation is positively correlated with genome size, suggesting insufficient time to purge TE insertions after a surge of activity. It is negatively correlated with TE diversity, suggesting genomic control mechanisms may limit the repertoire of active TEs ^30^. For specific TE classes, hotspots of accumulation dot the placental tree. For example, long interspersed nuclear elements (LINEs) are common in aardvarks and long terminal repeat (LTR) retrotransposons have accumulated in some rodents. Bats are a hotspot for horizontal transfer of DNA transposons (detailed in ^31^), having experienced over 200 such events, whereas only 11 transposons horizontally transferred into other lineages(**Table S4**).

### Evolutionary constraint in humans

Zoonomia was designed to achieve single-base resolution of constraint in placental mammals by maximizing branch length^16^. With the 240 species selected, only 191 positions in the human genome (0.000006%) are expected to be perfectly conserved by chance. In the Cactus alignment, we observe 19,000-fold more than our expectation (3.6 million perfectly conserved positions). We estimate that at least 10.7% (332 Mb) of the human genome is under constraint via purifying selection(**Fig. 2A**)^5^. While higher than estimated with the less well-powered 29 mammals dataset (5.5%)^32^, this is consistent with other estimates^33–35^(**Table S1**). We referenced the alignment on four other species (chimpanzee, mouse, dog and little brown bat) and measured constraint in each at different false discovery rates.

**Figure 2:**
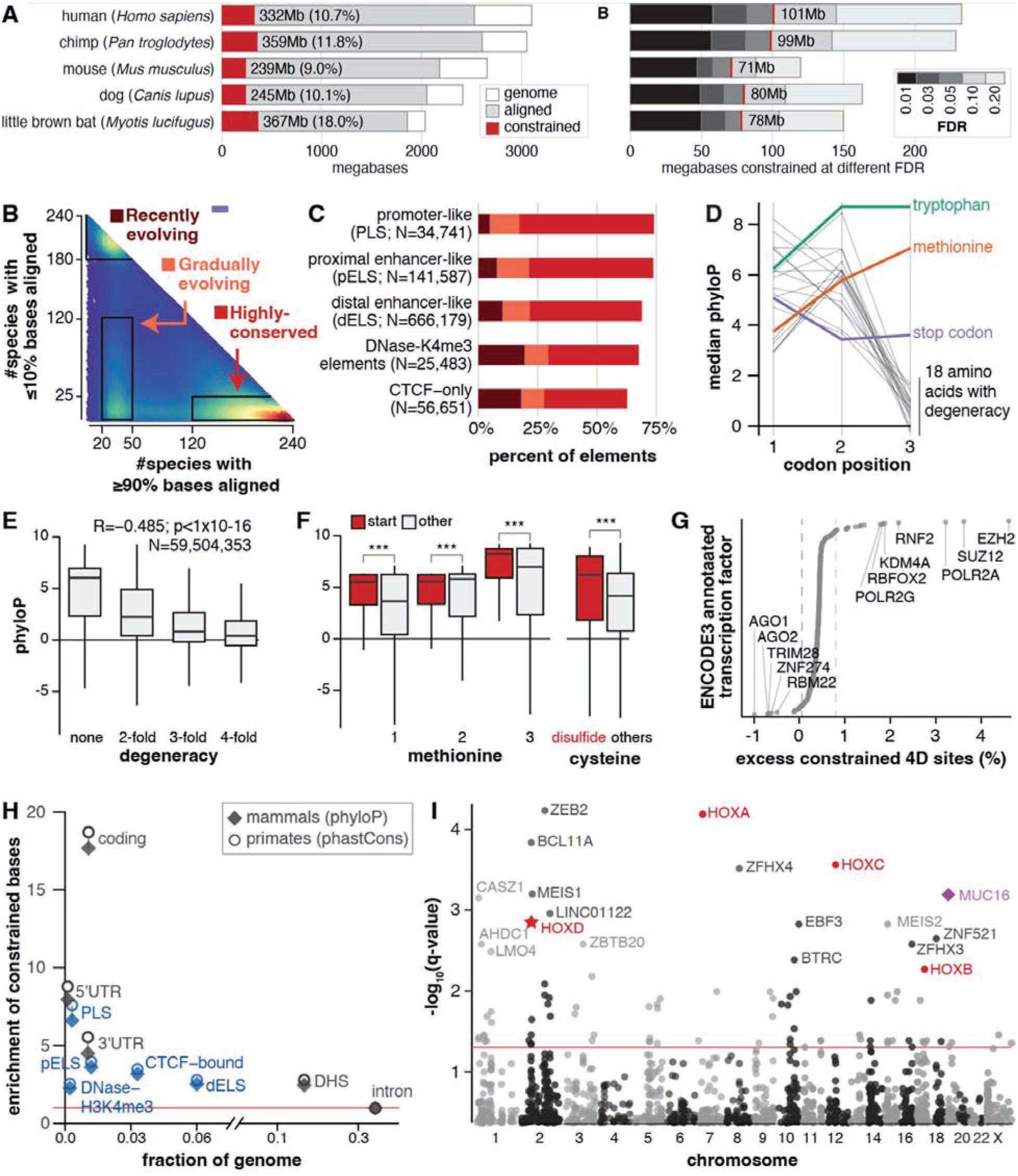
Comparing 240 species resolves mammalian constraint to single bases. **(A)** We estimated a lower-bound on the proportion of the genome under purifying selection (constraint) by comparing genome-wide phyloP to putatively neutrally evolving ancestral repeats and identified single bases as significantly constrained at varying FDR thresholds. (**B)** At 924,641 human cCREs^38^, we compared the number of species with poor alignments (y-axis) to the number with good alignments (x-axis) and found three clusters that are (**C)** non-randomly distributed across cCRE types. (**D)** Non-degenerate codons have higher phyloP scores. (**E)** Non-degenerate coding positions are most constrained; phyloP scores fall with increasing degeneracy. (**F)** Coding positions are more constrained if they function as start sites or form disulfide bridges (***p_Wilcoxon_<1×10^−16^). In E and F, the box represents 25%/75% quartiles, the horizontal line the median, and the vertical line the 5-95% quartiles. **(G)** Coding positions that are TFBS for some transcription factors are enriched for constraint. Dotted grey lines=5th and 95th percentiles. **(H)** Functional regions are enriched for constraint. cCREs are in blue. Red line: no enrichment. **(I)** The most constrained 100kb bins include all four HOX clusters (red text), with *HOXD* (star) overlapping the longest synteny block shared across mammals^51^. One bin (purple diamond) significantly lacks constraint and contains *MUC16*, a mucin secreted in innate immune responses. Red line: q=0.05. Bins with q< 0.006 are labelled.

We used the chimpanzee-referenced alignment to measure constraint at 10,032 human-specific deletions that had been functionally assessed using massively parallel reporter assays, including 717 associated with differences in enhancer activity (detailed in ^36^). Because these positions are deleted in humans, the human-referenced alignment does not score them. Filtering based on phyloP scores from the Cactus alignment substantially increased concordance between predicted transcription factor binding differences and measured species-specific regulatory activity.

#### Alignment depth

At any position in the 241-way Cactus alignment, the number of species aligned varies widely. Eleven percent of the human genome is aligned to ≥95% (≥228) of the species. Most of the genome (91%) aligns to at least five species (there are five other hominids in the alignment (**Fig. S2**). Highly conserved elements have deep alignments, and those evolving quickly align only to closely related species(**Fig. 2B**) (detailed in ^37^). In regulatory sequence, the depth of the alignment varies with element type. ENCODE^38^ promoter-like and enhancer-like candidate

*cis*-regulatory elements (cCREs) align well to most species, but CTCF-only cCREs tend to change more quickly(**Fig. 2C**).

#### Single-base resolution of constraint

Single bases confidently (FDR<0.05) designated as evolutionarily constrained comprise 3.53% (101 Mb) of the human genome (**Fig. 2A**; **Table S1**). In the human genome, most (80%) are within 5bp of another constrained base, and 30% are in blocks ≥5bp. Sequence data for ∼140,000 individuals in TOPMed^39^ shows variants at constrained bases (N=20,718,868) tend to have lower minor allele frequencies than unconstrained bases (0.0026 vs. 0.004; Wilcoxon rank sum exact test p = 9.5×10^−13^), consistent with purifying selection. Variants predicted to impact function have extremely low frequencies, regardless of phyloP score(**Fig. S3**).

Patterns of constraint in protein-coding sequence highlight the single-base resolution offered by the 241-way alignment. Most non-degenerate sites are constrained; most four-fold degenerate sites are not (74.1% vs 18.5%). Start codons, stop codons and splice sites are enriched for constraint (24x, 19x, and 25x greater; χ^2^ test, p < 2.2×10^−16^). For coding bases, median phyloP is 4.9 (IQR=5.8) in the first position (non-degenerate for 17/20 amino acids), rises to 6.0 (IQR=4.0) in the second (non-degenerate in 19/20), and drops to 0.68 (IQR=2.7) in the third (non-degenerate for 2/20)(**Fig. S4**). Stop codons and non-degenerate codons (tryptophan and methionine) are constrained at all three positions(**Fig. 2D**). PhyloP scores are negatively correlated with the number of nucleotide options (Spearman rho = −0.51, p < 2.2×10^−16^)(**Fig. 2E**). When codons encode amino acids at critical positions in the peptide, the variation in single-base constraint deviates from the pattern predicted from degeneracy alone. The codon for methionine is more conserved when functioning as a start codon than elsewhere(**Fig. 2F**). Similarly, cysteines in intra-peptide disulfide bridges (could cause misfolding when mutated) are more conserved than other cysteines(**Fig. 2F**). Four-fold degenerate sites overlapping ENCODE3 transcription factor binding sites (N = 2,647,541)^40^ are more constrained than other four-fold degenerate sites (N = 2,420,610; χ^2^ test, p<2.2×10^−16^, **Fig. 2G**, **Fig. S5**).

Constraint is strongly enriched in coding sequence and moderately enriched in regulatory elements(**Fig. 2H**). The most constrained genes are cell cycle and developmental, including embryology, neurogenesis, and morphogenesis of most organ systems. The least constrained genes shape an animal’s interaction with its environment: immunity (particularly response to pathogens), skin, smell, and taste(detailed in ^17^). Gene deserts (the longest 5% of intergenic regions; N=873) are more constrained when neighboring developmental transcription factors (N=224; p_Wilcoxon_=2.16×10^−15^). cCREs in these gene deserts (N=38,065) are unusually conserved(p_Wilcoxon_ < 2.2×10^−16^).

We used convolutional neural networks and publicly available ChIP-seq data to identify transcription factor binding sites (TFBS) genome-wide for 368 transcription factors(detailed in ^37^). Most TFBS are unconstrained, but for each transcription factor, a distinct minority are constrained (**Fig. 3A**) and appear active in other species (H3K4me3 and H3K27ac modified; **Fig. S6**). Motif scores for bases in constrained TFBS correlate with the magnitude of constraint(**Fig. 3BC**). Motif logos calculated from constrained TFBS for CTCF, a highly conserved and ubiquitously expressed transcription factor, are nearly identical across species(**Fig. 3D**). Most human CTCF constrained TFBS sites are found in macaque, rat, mouse and dog, and their function, as measured by CTCF ChiP-seq, is retained(**Fig. 3EF**). Unconstrained TFBS are often shared with macaque but not with more distantly related species.

**Figure 3:**
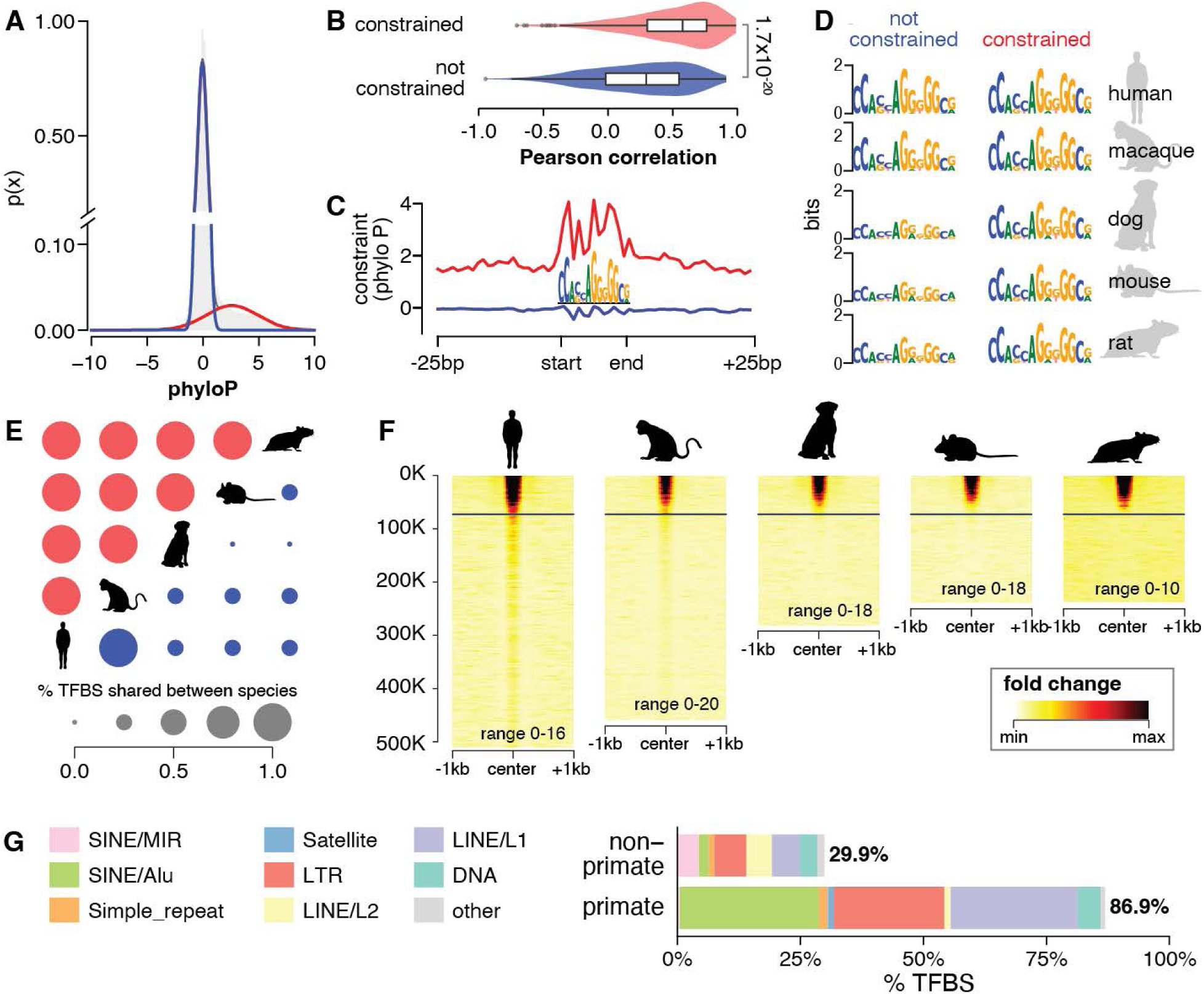
Conserved function of constrained CTCF transcription factor binding sites. (**A)** A two-component gaussian mixture model fit over average phyloP across binding sites for CTCF distinguishes distribution for evolutionarily constrained TFBS (red) and others (blue). (**B)** Across all transcription factors, aggregate phyloP is more correlated with TFBS information content for constrained than for non-constrained sites. (**C)** For example, at CTCF TFBS, aggregate phyloP scores for constrained (red) TFBS are high, but not for non-constrained (blue) TFBS (**D)** CTCF logos of constrained and non-constrained sets for four species made by lifting-over human TFBS. (**E)** Fraction of constrained (red) and non-constrained (blue) CTCF TFBSs shared between pairs of species. (**F)** CTCF transcription factor ChiP-seq signal over TFBSs sorted by average phyloP. Horizontal line indicates significant constraint.(**G)** Percent primate-specific and non-primate-specific TFBS that are derived from individual transposable element classes.

#### Regions of constraint

We defined constrained regions genome-wide using three approaches of varying stringency.

Zoonomia UltraConserved Elements (zooUCEs) are regions ≥20bp where every position is identical in at least 235 (98%) of species. This refines and expands the original set of 481 UCEs: elements >200 bp with identical sequence between human, mouse, and rat^41^. We discover 3,799 new UCEs, and 753 that overlap 318 original UCEs, for a total of 4,552 high-resolution zooUCEs averaging 28.9±13.0 bp, with 27 >= 100bp (**Fig. S7A**). ZooUCEs tend to be near genes in transcription-related and developmental biological processes(**Data S1**). The longest two (190bp and 161bp) are both in *POLA1,* which encodes the catalytic subunit of DNA polymerase-alpha, and are separated by a single base.

Regions of contiguous constraint (RoCC) are defined as twenty or more constrained bases in a row(**Fig. S7B**). This criteria, less stringent than for zooUCEs, finds 273 RoCCs ≥500bp and six ≥1kb. The longest RoCC (1.36kb; chr2:172071926-172073285) encompasses four distal enhancer-like ENCODE cCREs in an intron of the gene *METAP1D*. *METAP1D* encodes an essential mitochondrial protein conserved at least back to zebrafish^42^.

Finally, we divided the genome into 100kb bins (N=28,218) and identified 53 bins with significant constraint (q<0.05; average 17.8% constrained bases vs 3.5% for genome; **Table S5**). These bins are enriched for transcription-related biological processes (**Data S1**) and overlap all four *HOX* gene clusters (**Fig. 2I**). Five are in gene deserts, and two neighbor conserved developmental transcription factors (*LMO4* and *BCL11A*).

#### Unannotated constraint

Almost half of all constrained bases (48.5%) are in regions with no annotations in the thousands of cell types, tissues, or conditions assayed by ENCODE3 (**Table S6**)^38^. We grouped constrained bases <5bp apart in unannotated regions (excluding repeats, centromeres and telomeres) to define 424,179 <u>UN</u>annotated Intergenic <u>CO</u>nstrained <u>R</u>egio<u>N</u>s (UNICORNs) (mean size 38±53bp; Q95=131bp; 0.5% of genome; **Figs. 4A**, **S7C**). Most (77.0%) UNICORNs are within 500kb of the transcription start site for a protein-coding gene. UNICORNs have less variation than expected (lower minor allele frequencies; p_Wilcoxon_<2.2 x 10^−16^) and are less likely to overlap TOPMed^39^ variants(p_Wilcoxon_<2.2 x 10^−16^)(**Fig. 4B**).

**Figure 4:**
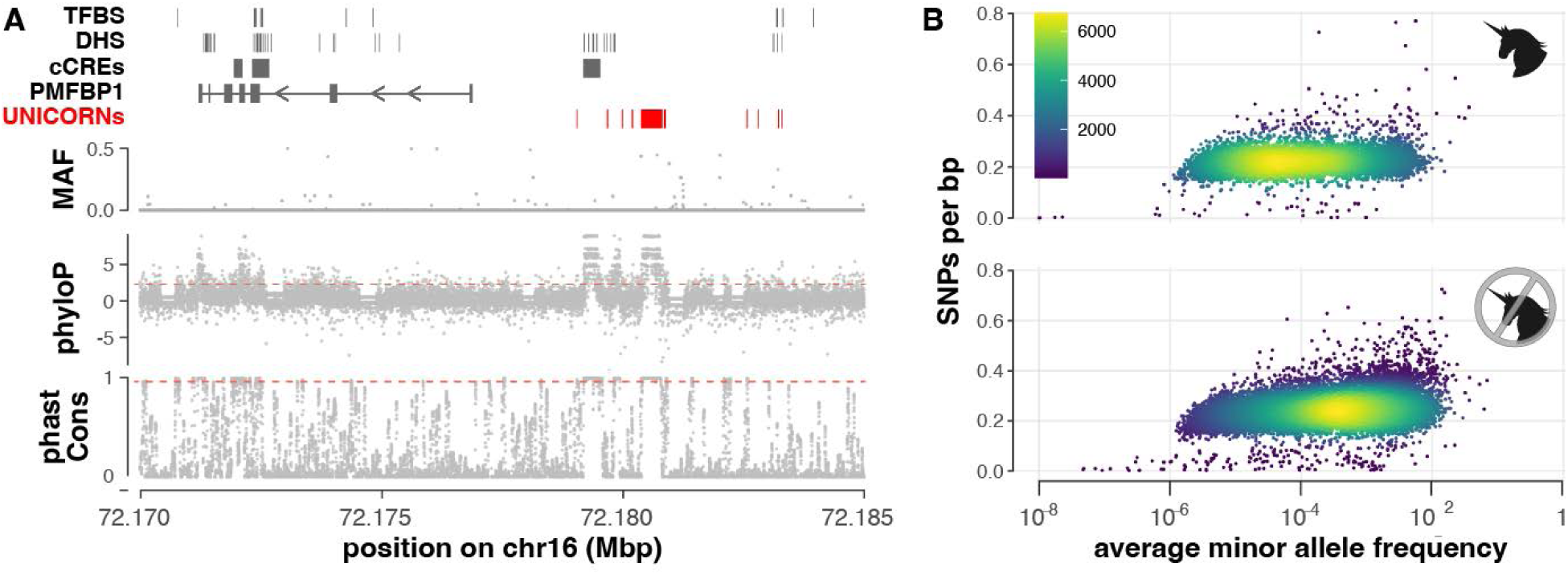
Constraint highlights unannotated regions that are likely functional. (**A)** Example UNICORNs (red) on human chromosome 16, the largest of which is 418 bp and located 3.5kb upstream of *PMFBP1*’s transcription start site. Bottom two tracks: mammal phyloP scores and primate phastCons scores. Grey dots represent single bases. Red dotted lines represent 5% FDR threshold for phyloP (phyloP > 2.270) and threshold for phastCons capturing equivalent genome proportion (phastCons base score ≥ 0.961). UNICORNs lack coding or regulatory annotations (top track) and have low diversity in human populations (second track and B). (**B)** Number of SNPs per base pair (0-1) against mean minor allele frequency for human SNPs within UNICORNs (top) or within a random set of unannotated sequences(bottom). Allele frequencies were log_10_ transformed. UNICORNs contain fewer variants, and those have lower allele frequencies than in the random set (Wilcoxon rank sum test, p < 2.2 x 10^−16^). Human variants and allele frequencies obtained from TOPMed data freeze 8^39^.

Many UNICORNs are likely functional under conditions not assayed in ENCODE3. Genomic elements that are active (annotated as open chromatin) in developing brain tissues^43^, adult motor cortical neuron cell types^44^, and narrowly defined regions of young adult brain^45^ overlap 8.8%, 7.1%, and 8.6% of UNICORNs, respectively (17% collectively). This suggests UNICORNs demarcate elements important in tissue types or at time points that are difficult to access in functional analyses.

#### Regulatory evolution through transposable elements

Overall, about 11% of constrained bases are in repeats in the human alignment. Constraint is enriched in simple repeats and DNA transposons and depleted in short interspersed nuclear elements, LTRs, and satellite repeats (**Fig. S8A**). This likely reflects differences in functional importance. Simple repeats closer to genes, and thus more likely to influence gene expression^46^, are more constrained (Spearman rho = −0.13, p < 2.2×10^−16^; **Fig. S8B**). DNA transposons are an ancient class of repeats known to acquire functional roles^47^, such as the transcription factor *ZBED5 (*70% constrained)^48^. In contrast, the repeat classes depleted in constraint have been active more recently during primate evolution and may not yet have acquired function^49^.

Most (87%) primate-specific TFBS overlap transposable elements (compared to 30% of non-primate TFBS), indicative of the important role of transposable elements in recent regulatory evolution^50^(**Fig. 3G**)(detailed in ^37^). TFBS in transposable elements, and especially younger elements, tend to be less conserved and change more quickly(**Fig. S9A,B**)

#### Accelerated regions and topological associating domain boundaries

We developed an automated pipeline for identifying “accelerated” regions that are highly constrained across placental mammals but exceptionally variable in particular lineages, suggesting adaptive evolution (detailed in ^51^). Applying this to the human and chimp alignments, we found 312 human accelerated regions (zooHARs) and 141 in chimpanzees (zooCHARs). Accelerated regions in both species are mainly noncoding, have signatures of positive selection (82% and 86%, respectively), and reside near developmental and neurological genes^52^, like previous sets of HARs^53–55^. zooHARs and zooCHARs overlap topological associating domains containing human-specific and chimp-specific structural variants, respectively^56^. 3D genome structure is altered by human-specific structural variants in zooHARs, suggesting a mechanism for species-specific evolution through enhancer hijacking^51^.

### Annotation and alignment of protein-coding genes

#### Annotating genes through alignment

Pairing the Cactus alignment with gene annotations from three species with exceptionally high-quality, high-contiguity genomes (human, mouse, and goat), we annotated genes in 238 placental species, including 129 without an NCBI or Ensembl annotation (**Table S2**)^57^. Predicted orthologs have contiguous open reading frames spanning 90-110% of human protein length. Starting with an input set of 22,799 protein-coding genes, we annotated between 3,871 and 20,168 (averaging 12,905 ± 2,925) genes per genome. As expected, the number of genes annotated decreases as assembly contiguity drops (contig N50: Pearson r=0.28, p=1×10^−5^; scaffold N50: Pearson r=0.37, p=3×10^−9^) and as phylogenetic distance from human (the species with the highest quality gene annotations) increases (Pearson r=-0.41, p=7×10^−11^) (**Table S7**). These annotations, although incomplete, offer the first gene annotations for over a hundred species.

#### New 428-way protein-coding alignment in mammals

We constructed the largest-ever mammalian protein-coding alignment (428 species) using a new method integrating gene annotation, ortholog detection, and classification of genes as intact or lost (detailed in ^18^). TOGA (Tool to infer Orthologs from Genome Alignments) uses a machine learning model trained on human and mouse to annotate orthologs between human and each of the other species, and it can distinguish orthologs from paralogs or processed pseudogenes. It joins non-contiguous genes in fragmented assemblies, making it particularly useful for species where obtaining high-quality DNA is difficult, including rare or endangered species. We annotated, on average, 19,144 orthologous genes for ape assemblies and 17,779 genes for non-ape assemblies. Using a benchmarking set of mammalian single copy orthologs^58^, TOGA improves annotation completeness for 108 of the 118 assemblies with an existing NCBI annotation.

#### Repeated loss of CMAH gene is associated with immunity evolution

For one gene known to be lost in humans, we used non-human referenced alignments to explore its persistence across placental mammals. Loss of *CMAH* was initially thought to be specific to the *Homo* lineage, prompting speculation it might explain distinctive human characteristics^59–62^. CMAH hydroxylates the sialic acid Neu5Ac to Neu5Gc, and its loss restricts infection by pathogens dependent on Neu5Gc (e.g. *Plasmodium reichenowi*^63^) but increases susceptibility to viruses that bind Neu5Ac (e.g. MERS coronavirus^64^) and may drive speciation^65^. We found three independent losses of *CMAH* in rodents and three in bats (two already described^62^)(**Fig. S10**). *CMAH* accumulated multiple stop codons in the Cairo spiny mouse. Loss in mountain beaver and Brazilian guinea pig reflects diminished gene content and pseudogenization. Additional experiments are needed to confirm non-functionality.

Among the species with *CMAH* loss is the American mink (*Neovison vison),* the only non-human animal to suffer well-documented high mortality from SARS-CoV2. SARS-CoV2 likely lacks Neu5Gc^66^ and thus may face fewer challenges in moving among mammalian hosts lacking *CMAH* ^67, 68^. *CMAH* loss has been described in white-tailed deer^62^, another species susceptible to SARS-CoV-2 infection (without high mortality)^69^, but we found fully intact *CMAH* homologs in more recent assemblies (GCA_002102435.1 and GCA_014726795.1). The connection between *CMAH* loss and zoonotic potential warrants further investigation, and monitoring species that have lost *CMAH* may help anticipate future zoonotic transfer events^70^.

### Connecting genotype to phenotype

#### Olfaction

We performed *de novo* olfactory receptor gene detection in 249 species and confirmed that counts vary widely (mean count = 1,217±683; N=249)(**Table S8**)^71^. Our reference-free approach is designed to avoid underestimating olfactory receptor gene counts in distantly related taxa and is robust to variation in genome assembly quality (count ∼ contig N50: rho = 0.006, p-value= 0.93; count ∼ scaffold N50: rho = 0.013; p-value= 0.84; N=236). African elephant has the largest repertoire (4,199 genes; 1,765 functional and 2,434 non-functional); killer whale has the smallest (154 genes: 24 functional and 130 non-functional)(**Fig. S11**).

Similar olfactory receptor gene counts in distantly related species may reflect shared selection pressures. Bumblebee bat and North Pacific right whale have among the smallest repertoires (332 and 392 genes respectively). Both species rely on echolocation ^72^, although North Pacific right whale is also aquatic. Overall, whales (17 species) have fewer than any other clade, averaging 225±75 genes, consistent with previous work^73^. Closely related species can have divergent counts. Anteaters, tree sloths, and armadillos (superorder Xenarthra) generally have large repertoires (2,784±1,089 genes, N=8). Brown-throated three-toed sloth, though, has only 545 genes (115 functional, 430 non-functional), far fewer than the two-toed sloth (2,314 genes: 929 functional, 1,385 non-functional) and Hoffman’s two-toed sloth (3,839 genes, 1,268 functional, 2,571 non-functional). Three-toed sloths (family Bradypodidae) are less active and have better vision than two-toed sloths (family Choloepodidae), which rely on olfaction to explore new locations^74^. The olfactory variation between the two sloth families supports the hypothesis that they independently evolved suspensory behavior^75–77^.

Olfactory receptor gene counts are strongly correlated with olfactory turbinal counts(Spearman rho=0.71; phylolm coefficient 0.014, phylogenetic permutations p = 0.0013; N=63)(**Fig. S12**) ^78, 79^. Turbinals are an anatomic feature of the nasal cavity associated with olfaction capacity ^80–82^. With 65 phenotyped species, we discerned the connection between these two traits through genomic analysis, highlighting the potential of this approach for investigating multifactorial evolutionary responses to a selective pressure.

#### Hibernation

The evolutionary origins of hibernation and its core physiological state of metabolic depression (torpor) are unclear. Hibernation exists in every deep mammalian lineage, suggesting that it is either an ancestral trait repeatedly lost or a derived trait repeatedly gained^83, 84^. Understanding the genomic basis of its constituent physiology, including cellular recovery from repeated cooling and rewarming without apparent long-term harm, could inform therapeutics, critical care, and long-distance spaceflight.

If hibernation is ancestral, hibernator orthologs should more closely match the mammalian ancestor, revealing putative essential genes underlying this phenotype. We used Generalized Least Squares Forward Genomics^13^ to identify 100bp protein-coding regions where 22 deep hibernators (species capable of core temperature depression below 18°C for >24h) had higher sequence identity than 154 strict homeotherms to a reconstructed ancestral mammal genome. We found 14 regions in 11 genes more conserved (p_FD R_ <0.01) in deep hibernators than in strict homeotherms. Two genes, *MFN2* and *PINK1*, overlap four gene ontology (GO) Biological Process gene sets (p_FDR_<0.05) related to depolarization and degradation of damaged mitochondria, an organelle essential for metabolic depression^85^. A third, *TXNIP,* also regulates mitophagy^86^, with torpor-responsive gene expression occurring in several lineages of hibernators ^87–89^(**Table S9**).

We did a second, distinct analysis of hibernation by applying RERconverge^90^ to our TOGA alignments. We detected 764 genes with disparate evolutionary rates in 21 deep hibernators versus 146 non-hibernators(p_FDR_<0.05; **Data S3**, **Fig. S13**). 511 genes score as slower-evolving in hibernators, and 253 as more quickly evolving. The duration and depth of hibernation are highly variable across mammals, suggesting evolutionary honing of the hibernation phenotype in different lineages. Consistent with this, both fast and slow evolving genes overlap 6,505 genes identified as torpor-modified in hibernator brain^87^

Genes evolving more quickly in hibernators are enriched in GO pathways relating to synaptic transmission; neuroprotection in the cold is well-documented for hibernator brains, as is their ability to rapidly remodel synapses upon rewarming^91^. Genes evolving more slowly in hibernators are enriched in GO pathways related to DNA repair, suggesting that an effective response to DNA damage may be essential to successful hibernation^92^. In both gene lists, metabolic processes are an over-represented pathway, consistent with the importance of metabolic regulation in hibernation^85, 93^.

#### Regulatory evolution

We developed a forward genomics toolkit for associating *cis*-regulatory elements with specific phenotypes (detailed in ^94^). This Tissue-Aware Conservation Inference Toolkit (TACIT) uses open chromatin sequence data available for a few species to predict open chromatin (a proxy for *cis*-regulatory element activity) in many species at open chromatin region orthologs found by the Cactus alignment(**Fig. 5A**)^95^. TACIT then associates open chromatin predictions with phenotype annotations. By using open chromatin predictions, TACIT does not require tissue-specific open chromatin data in each species, which is costly and logistically challenging to obtain.

**Figure 5:**
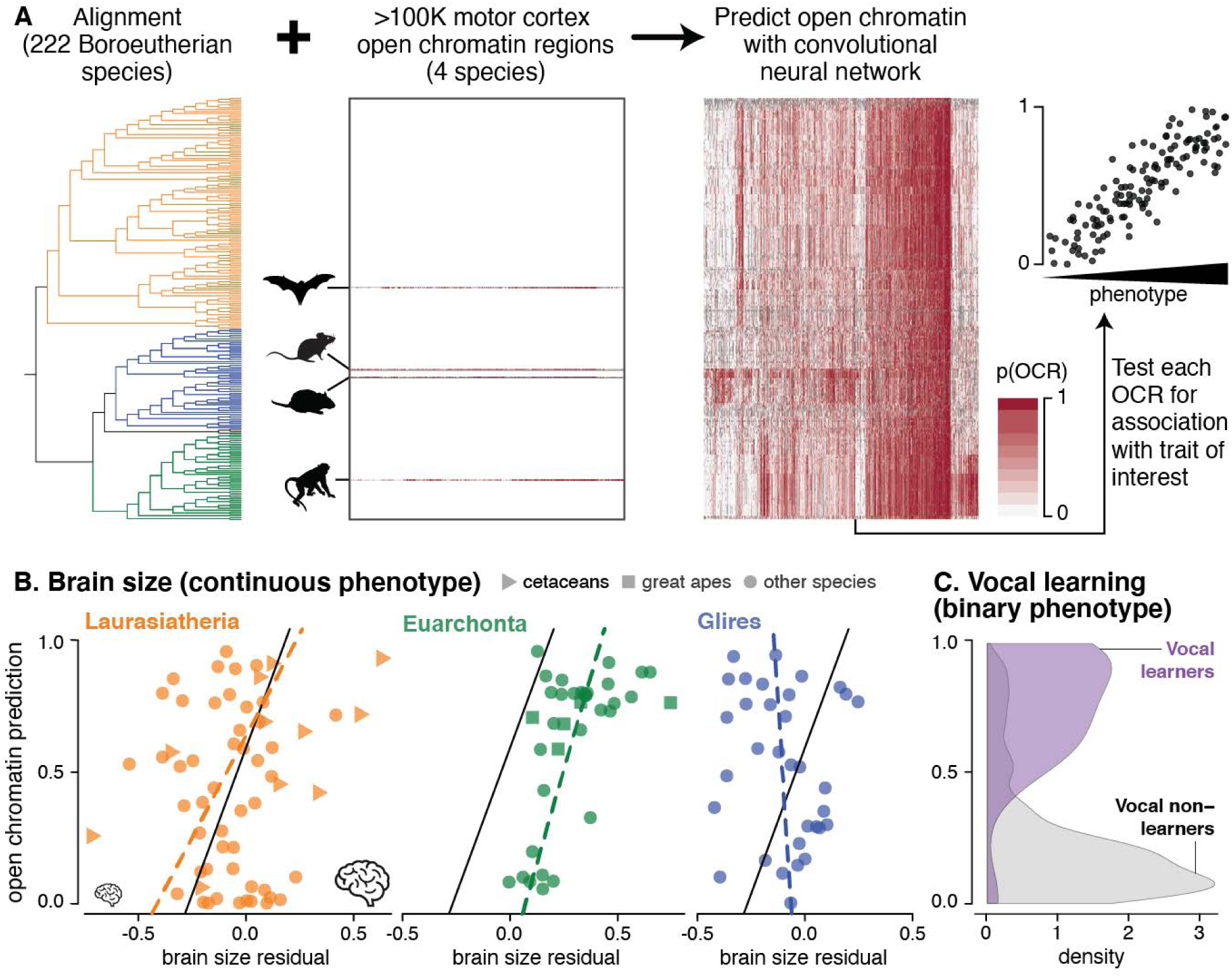
Forward genomics associates genomic differences with brain size and vocal learning. **(A)** TACIT trains a convolutional neural network on open chromatin regions in a few species and then predicts open chromatin in specific tissues or cell types in many others and screens for phenotype associations. **(B)** TACIT associates motor cortex open chromatin with brain size (a quantitative trait), and here we show results for the rhesus macaque open chromatin region chr10:48660711-48661679 in the *MACROD2* locus, which is implicated in nervous system development^106^. The solid line shows the phylolm line of best fit (coeff. = 0.451, permulations p_FDR_= 0.0375). The dashed lines show phylolm lines of best fit for each clade as a visual aid. **(C)** We also applied TACIT to associate motor cortex open chromatin with vocal learning, a binary trait, and here we show results for the Egyptian fruit bat open chromatin region PVIL01002568.1:139004-139596 in the *GALC* locus (permulations p_FDR_=0.0376), which was also associated with open chromatin specifically active in the orofacial motor cortex of bat^97^ and which has previously been associated with severe speech and motor disabilities in humans^99^.

We applied TACIT to neural traits with substantial species-level phenotype data: brain size (158 species) and vocal learning (181 species). We first predicted open chromatin using models trained on data from different parts of the brain and then tested for associations. For brain size, we found 34 motor cortex (**Fig. 5B**) and 13 parvalbumin neuron associations. Motor cortex open chromatin regions near microcephaly or macrocephaly genes had stronger associations (1-sided p_Wilcoxon_=0.0073).

For vocal learning, we tested for both regulatory and coding changes using TACIT and RERconverge (detailed in ^97^). Vocal learning is the ability to mimic non-innate sounds, and it likely evolved convergently in humans, bats, cetaceans, and pinnipeds^98^. Using TACIT, we associated open chromatin regions near *GALC*^99^(**Fig. 5C**) and other speech disorder-related genes with vocal learning. This included a region near *TSHZ3,* a transcription factor whose disruption results in absence or delay of human speech^100^.

### Biodiversity

We built a new tool to identify species in urgent need of conservation using genomic features correlated to elevated risk of extinction (detailed in ^101^). While no single genomic summary statistic is diagnostic of conservation classification by the International Union for Conservation of Nature (IUCN), threatened species have, on average, lower heterozygosity (phylogenetic regression p=0.016)^16^, smaller long-term effective population sizes (N_e_)(p<3.30×10^−5^), larger recent declines in population size (ratio of historic N_e_ to contemporary population size; p=0.012), higher proportions of missense mutations (p=3.63×10^−4^), and fewer mutations at extremely conserved sites, as measured by kurtosis of the phyloP distribution(p=0.004; **Fig. 6**). The predictive value of N_e_ is especially striking given that historic N_e_ reflects population dynamics more than 10,000 years ago. Both long-term population history and contemporary declines appear to imperil species survival^101^.

**Figure 6.**
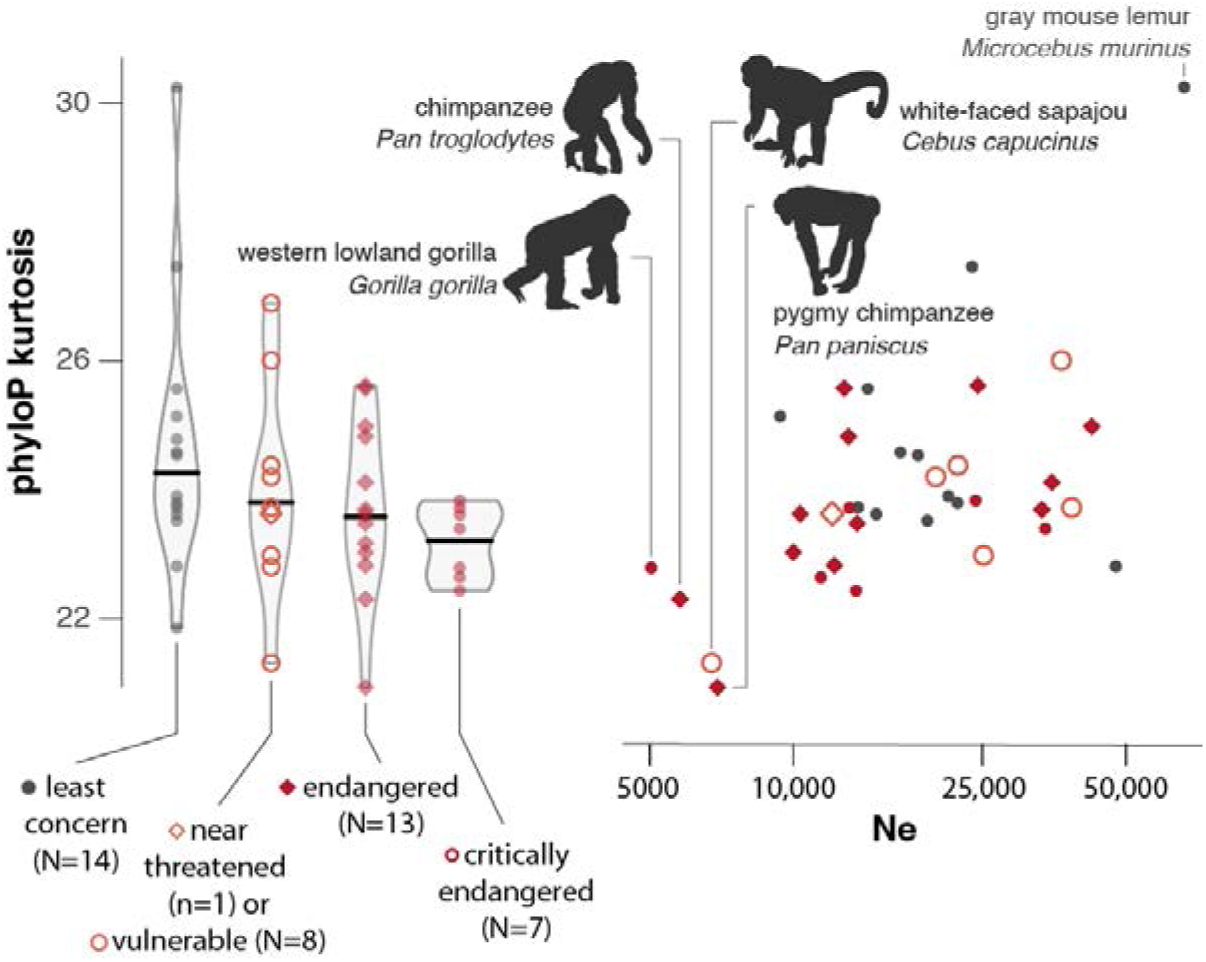
Genomic metrics distinguish at-risk species. Primates categorized at increasing levels of extinction risk tend to have fewer mutations at sites with extreme phyloP scores, measured as phyloP kurtosis, and smaller effective population sizes (*N_e_*). The four species (all at risk) with the lowest *N_e_* and the species with the highest *N_e_* are labeled. Screening several genome metrics in parallel provides initial assessment of extinction risk that can help prioritize species for deeper assessment ^101^.

Applying predictive models using genomic metrics to species lacking IUCN classifications, we assess the Upper Galilee Mountains blind mole rat as unlikely to be threatened (12-14% probability). The risk is higher for Java lesser chevrotain (27-63%) and for killer whale (53-69%), with several populations already listed as endangered^102^. Our findings suggest a single reference genome, an increasingly achievable resource^103^, may be valuable for identifying species in need of further study and immediate intervention.

## Conclusion

In 2011, the landmark 29 Mammals paper demonstrated that evolutionary constraint is a powerful indicator of function^32^ and estimated that sequencing a few hundred placental mammals would achieve single-nucleotide resolution of constraint across the human genome. Zoonomia realizes that vision. In combination with existing resources such as ENCODE^38^ and TOPMed^39^ and new technologies including regulatory genomic assays^94, 97^, chromatin conformation capture^27, 51^, and massively parallel reporter assays^36^, Zoonomia’s alignments of hundreds of placental species address fundamental questions in mammalian evolution.

Zoonomia addresses the central goal of medical genomics: to identify genetic variants that influence disease risk and understand their biological mechanisms^104^. Constraint scores confirm strong conservation of protein-coding sequences and discern subtle correlation to other functions, like transcription factor binding. Unsurprisingly, *HOX* gene clusters are exceptionally constrained. More surprisingly, constraint is nearly as strong in gene deserts neighboring developmental genes. Constrained regions overall, including those from Zoonomia’s updated list of ultraconserved elements and new list of UNICORNs, are strongly enriched for transcriptional regulation and developmental genes. Even in regions of only modest nucleotide-level conservation, our alignments help predict activity conservation at orthologs via prediction of tissue and cell type-specific *cis*-regulatory elements across species. Hundreds of other regions, mainly non-coding, are accelerated, especially in primates and humans^51^. They are enriched for topologically associated domains containing human-specific structural variants, perhaps reflecting adaptation via altered 3D genome structure.

Zoonomia marks a new era in non-human and comparative genomics while underscoring the critical need for high-quality mammalian phenotypes. Our alignments facilitate annotation of coding, regulatory, and functionally constrained regions in hundreds of species. Forward genomic approaches can find genes and regulatory elements underlying traits that, while medically relevant, are difficult to study in humans because our species is either not exceptional (e.g., hibernation and olfaction) or has minimal natural variation (e.g., brain size and vocal learning). Using just the 27% of Zoonomia species for which anatomic turbinal-count data are available, we discovered a strong correlation with olfactory receptor gene counts. Even so, achieving the richer data sets needed to study other phenotypes, evaluate pattern robustness, and address broader prospects — for example, the possibility of predicting genomic features from fossils^9^ — will require close collaborations between genomics researchers and scientists with expertise in morphology, physiology, and behavior to prioritize species for paired sequencing and phenotyping.

Comparative genomics projects are classically motivated by the potential to advance human biomedicine, all while relying on biodiversity imperiled by human activity^105^. We show that even a single reference genome per species, the core data type for comparative genomics, can support biodiversity conservation by guiding extinction risk assessment^101^. With genome assemblies increasingly accessible^103^, conservation scientists will be better equipped to identify potentially threatened populations earlier, when management efforts are more effective.

Through close and enduring partnerships among researchers in genomics, morphology, behavior, and conservation, resources from Zoonomia and other comparative-genomics projects can simultaneously address questions in human health and trait mapping while guiding conservation of the biodiversity that is essential to these discoveries.

## Acknowledgments

We thank everyone who collected the samples and phenotype data essential for this project. We especially thank the San Diego Zoo Wildlife Alliance staff, E. Baitchman. R. Johnston, and Zoo New England, Broad Institute Genomics Platform, the SNP & SEQ Technology Platform (National Genomics Infrastructure Sweden and Science for Life Laboratory), and the Texas A&M High Performance Research Computing Center. Species silhouettes were sourced from PhyloPic.

## Data and materials availability

Zoonomia data resources are available through the UCSC genome browser, NCBI, and github. Data and materials for companion papers can be found within the specific papers.

## Consortium list

Gregory Andrews^1^, Joel C. Armstrong^2^, Matteo Bianchi^3^, Bruce W. Birren^4^, Kevin R. Bredemeyer^5^, Ana M. Breit^6^, Matthew J. Christmas^3^, Hiram Clawson^2^, Joana Damas^7^, Federica Di Palma^8, 9^, Mark Diekhans^2^, Michael X. Dong^3^, Eduardo Eizirik^10^, Kaili Fan^1^, Cornelia Fanter^11^, Nicole M. Foley^5^, Karin Forsberg-Nilsson^12, 13^, Carlos J. Garcia^14^, John Gatesy^15^, Steven Gazal^16^, Diane P. Genereux^4^, Linda Goodman^17^, Jenna Grimshaw^14^, Michaela K. Halsey^14^, Andrew J. Harris^5^, Glenn Hickey^18^, Michael Hiller^19–21^, Allyson G. Hindle^11^, Robert M. Hubley^22^, Graham M. Hughes^23^, Jeremy Johnson^4^, David Juan^24^, Irene M. Kaplow^25, 26^, Elinor K. Karlsson^1, 4, 27^, Kathleen C. Keough^17, 28, 29^, Bogdan Kirilenko^19–21^, Klaus-Peter Koepfli^30–32^, Jennifer M. Korstian^14^, Amanda Kowalczyk^25, 26^, Sergey V. Kozyrev^3^, Alyssa J. Lawler^4, 26, 33^, Colleen Lawless^23^, Thomas Lehmann^34^, Danielle L. Levesque^6^, Harris A. Lewin^7, 35, 36^, Xue Li^1, 4, 37^, Abigail Lind^38^, Kerstin Lindblad-Toh^3, 4^, Ava Mackay-Smith^39^, Voichita D. Marinescu^3^, Tomas Marques-Bonet^40–43^, Victor C. Mason^44^, Jennifer R. S. Meadows^3^, Wynn K. Meyer^45^, Jill E. Moore^1^, Lucas R. Moreira^1, 4^, Diana D. Moreno-Santillan^14^, Kathleen M. Morrill^1, 4, 37^, Gerard Muntané^24^, William J. Murphy^5^, Arcadi Navarro^40, 42, 46, 47^, Martin Nweeia^48–51^, Sylvia Ortmann^52^, Austin Osmanski^14^, Benedict Paten^2^, Nicole S. Paulat^14^, Andreas R. Pfenning^25, 26^, BaDoi N. Phan^25, 26, 53^, Katherine S. Pollard^28, 29, 54^, Henry E. Pratt^1^, David A. Ray^14^, Steven K. Reilly^39^, Jeb R. Rosen^22^, Irina Ruf^55^, Louise Ryan^23^, Oliver A. Ryder^56,57^, Pardis C. Sabeti^4,58,59^, Daniel E. Schäffer^25^, Aitor Serres^24^, Beth Shapiro^60,61^, Arian F. A. Smit^22^, Mark Springer^62^, Chaitanya Srinivasan^25^, Cynthia Steiner^56^, Jessica M. Storer^22^, Kevin A. M. Sullivan^14^, Patrick F. Sullivan^63,64^, Elisabeth Sundström^3^, Megan A. Supple^60^, Ross Swofford^4^, Joy-El Talbot^65^, Emma Teeling^23^, Jason Turner-Maier^4^, Alejandro Valenzuela^24^, Franziska Wagner^66^, Ola Wallerman^3^, Chao Wang^3^, Juehan Wang^16^, Zhiping Weng^1^, Aryn P. Wilder^56^, Morgan E. Wirthlin^25,26,67^, James R. Xue^4,58^, Xiaomeng Zhang^4,25,26^

Affiliations:

^1^Program in Bioinformatics and Integrative Biology, UMass Chan Medical School; Worcester, MA 01605, USA.

^2^Genomics Institute, University of California Santa Cruz; Santa Cruz, CA 95064, USA.

^3^Department of Medical Biochemistry and Microbiology, Science for Life Laboratory, Uppsala University; Uppsala, 751 32, Sweden.

^4^Broad Institute of MIT and Harvard; Cambridge, MA 02139, USA.

^5^Veterinary Integrative Biosciences, Texas A&M University; College Station, TX 77843, USA.

^6^School of Biology and Ecology, University of Maine; Orono, ME 04469, USA.

^7^The Genome Center, University of California Davis; Davis, CA 95616, USA.

^8^Genome British Columbia; Vancouver, BC, Canada.

^9^School of Biological Sciences, University of East Anglia; Norwich, UK.

^10^School of Health and Life Sciences, Pontifical Catholic University of Rio Grande do Sul; Porto Alegre, 90619-900, Brazil.

^11^School of Life Sciences, University of Nevada Las Vegas; Las Vegas, NV 89154, USA.

^12^Biodiscovery Institute, University of Nottingham; Nottingham, UK.

^13^Department of Immunology, Genetics and Pathology, Science for Life Laboratory, Uppsala University; Uppsala, 751 85, Sweden.

^14^Department of Biological Sciences, Texas Tech University; Lubbock, TX 79409, USA.

^15^Division of Vertebrate Zoology, American Museum of Natural History; New York, NY 10024, USA.

^16^Keck School of Medicine, University of Southern California; Los Angeles, CA 90033, USA.

^17^Fauna Bio Incorporated; Emeryville, CA 94608, USA.

^18^Baskin School of Engineering, University of California Santa Cruz; Santa Cruz, CA 95064, USA.

^19^Faculty of Biosciences, Goethe-University; 60438 Frankfurt, Germany.

^20^LOEWE Centre for Translational Biodiversity Genomics; 60325 Frankfurt, Germany.

^21^Senckenberg Research Institute; 60325 Frankfurt, Germany.

^22^Institute for Systems Biology; Seattle, WA 98109, USA.

^23^School of Biology and Environmental Science, University College Dublin; Belfield, Dublin 4, Ireland.

^24^Department of Experimental and Health Sciences, Institute of Evolutionary Biology (UPF-CSIC), Universitat Pompeu Fabra; Barcelona, 08003, Spain.

^25^Department of Computational Biology, School of Computer Science, Carnegie Mellon University; Pittsburgh, PA 15213, USA.

^26^Neuroscience Institute, Carnegie Mellon University; Pittsburgh, PA 15213, USA.

^27^Program in Molecular Medicine, UMass Chan Medical School; Worcester, MA 01605, USA.

^28^Department of Epidemiology & Biostatistics, University of California San Francisco; San Francisco, CA 94158, USA.

^29^Gladstone Institutes; San Francisco, CA 94158, USA.

^30^Center for Species Survival, Smithsonian’s National Zoo and Conservation Biology Institute; Washington, DC 20008, USA.

^31^Computer Technologies Laboratory, ITMO University; St. Petersburg 197101, Russia.

^32^Smithsonian-Mason School of Conservation, George Mason University; Front Royal, VA 22630, USA.

^33^Department of Biological Sciences, Mellon College of Science, Carnegie Mellon University; Pittsburgh, PA 15213, USA.

^34^Senckenberg Research Institute and Natural History Museum Frankfurt; 60325 Frankfurt am Main, Germany.

^35^Department of Evolution and Ecology, University of California Davis; Davis, CA 95616, USA.

^36^John Muir Institute for the Environment, University of California Davis; Davis, CA 95616, USA.

^37^Morningside Graduate School of Biomedical Sciences, UMass Chan Medical School; Worcester, MA 01605, USA.

^38^NA.

^39^Department of Genetics, Yale School of Medicine; New Haven, CT 06510, USA.

^40^Catalan Institution of Research and Advanced Studies (ICREA); Barcelona, 08010, Spain.

^41^CNAG-CRG, Centre for Genomic Regulation, Barcelona Institute of Science and Technology (BIST); Barcelona, 08036, Spain.

^42^Department of Medicine and LIfe Sciences, Institute of Evolutionary Biology (UPF-CSIC), Universitat Pompeu Fabra; Barcelona, 08003, Spain.

^43^Institut Catalàde Paleontologia Miquel Crusafont, Universitat Autònoma de Barcelona; 08193, Cerdanyola del Vallès, Barcelona, Spain.

^44^Institute of Cell Biology, University of Bern; 3012, Bern, Switzerland.

^45^Department of Biological Sciences, Lehigh University; Bethlehem, PA 18015, USA.

^46^BarcelonaBeta Brain Research Center, Pasqual Maragall Foundation; Barcelona, 08005, Spain.

^47^CRG, Centre for Genomic Regulation, Barcelona Institute of Science and Technology (BIST); Barcelona, 08003, Spain.

^48^Department of Comprehensive Care, School of Dental Medicine, Case Western Reserve University; Cleveland, OH 44106, USA.

^49^Department of Vertebrate Zoology, Canadian Museum of Nature; Ottawa, Ontario K2P 2R1, Canada.

^50^Department of Vertebrate Zoology, Smithsonian Institution; Washington, DC 20002, USA.

^51^Narwhal Genome Initiative, Department of Restorative Dentistry and Biomaterials Sciences, Harvard School of Dental Medicine; Boston, MA 02115, USA.

^52^Department of Evolutionary Ecology, Leibniz Institute for Zoo and Wildlife Research; 10315 Berlin, Germany.

^53^Medical Scientist Training Program, University of Pittsburgh School of Medicine; Pittsburgh, PA 15261, USA.

^54^Chan Zuckerberg Biohub; San Francisco, CA 94158, USA.

^55^Division of Messel Research and Mammalogy, Senckenberg Research Institute and Natural History Museum Frankfurt; 60325 Frankfurt am Main, Germany.

^56^Conservation Genetics, San Diego Zoo Wildlife Alliance; Escondido, CA 92027, USA.

^57^Department of Evolution, Behavior and Ecology, School of Biological Sciences, University of California San Diego; La Jolla, CA 92039, USA.

^58^Department of Organismic and Evolutionary Biology, Harvard University; Cambridge, MA 02138, USA.

^59^Howard Hughes Medical Institute; Chevy Chase, MD, USA.

^60^Department of Ecology and Evolutionary Biology, University of California Santa Cruz; Santa Cruz, CA 95064, USA.

^61^Howard Hughes Medical Institute, University of California Santa Cruz; Santa Cruz, CA 95064, USA.

^62^Department of Evolution, Ecology and Organismal Biology, University of California Riverside; Riverside, CA 92521, USA.

^63^Department of Genetics, University of North Carolina Medical School; Chapel Hill, NC 27599, USA.

^64^Department of Medical Epidemiology and Biostatistics, Karolinska Institutet; Stockholm, Sweden.

^65^Iris Data Solutions, LLC; Orono, ME 04473, USA.

^66^Museum of Zoology, Senckenberg Natural History Collections Dresden; 01109 Dresden, Germany.

^67^Allen Institute for Brain Science; Seattle, WA 98109, USA

