## Supplemental information for "Evolutionary constraint and innovation across hundreds of placental mammals"

Materials and Methods

Alignment update

After inferring the initial 242-way alignment used in(*14*), we discovered that the tarsier (*Carlito syrichta*) genome we had included was a mislabeled kangaroo rat. We removed the mislabeled genome from the alignment using the halRemoveGenome command in the HAL toolkit(*107*). But, due to the progressive alignment process, the misplaced genome likely negatively influenced the reconstruction of the ancestors above it in the guide tree and thus the alignment from the rest of the primates (the clade it was basal to) to the other clades. To remedy this, we re-aligned the affected part of the alignment (the parts involved in inferring all ancestors above tarsier) to remove the influence of the mislabeled genome without requiring re-alignment of unaffected parts of the tree. Since subtrees not below tarsier were not affected (due to their alignments not including tarsier as a descendant or outgroup genome), they did not require re-inference. We constructed a re-inference alignment with guide tree:

((((((fullTreeAnc110:0.1523,Galeopterus_variegatus:0.12765)fullTreeAnc112:0.01,fullTreeAnc113:0.15442)fullTreeAnc114:0.04,fullTreeAnc69:0.04)fullTreeAnc115:0.0406,fullTreeAnc237:0.0212)fullTreeAnc238:0.0237,fullTreeAnc14:0.02)fullTreeAnc239;

where we obtained the *fullTreeAnc* ancestors used as leaves from the original HAL. We used platypus as an outgroup genome. We used the relationships and ancestral sequences from the re-inferred "supertree" alignment and merged them with the existing, original alignments between species in each subtree (using halReplaceGenome from the HAL toolkit). In the process, we removed the now-redundant ancestor above tarsier (*fullTreeAnc111*). The alignment improved after removal of the duplicate genome and re-inference: coverage of the human genome from non-primate species increased by an average of 2.2% of the human genome, which was an average 6% relative increase from the coverage in the original alignment.

Curating transposable elements

We gathered final genome assemblies for each species under examination and identified putative TEs using RepeatModeler-4.0.9(*108*). We rejected some assemblies from the analysis for either of two reasons: 1) the presence of multiple clearly identifiable TE assembly artifacts (n=5), which decrease confidence in the accuracy of TE consensus estimates or 2) the availability of an existing, well-curated species-specific library from RepBase(*109*) or from efforts for other projects in our own laboratory or others (n=29; **Table S2**), which would make new analysis a duplication of efforts. Using the RepeatModeler output from each genome, we then embarked on a species-by-species curation of the putative elements using methods described in our previous work(*110*).

To minimize the potential for re-curating known elements, we presumed that only younger TEs, i.e. consensus sequences having hits with K2P distances(*111*) less than 4.4% (~20my or less since insertion) based on a general mammalian neutral mutation rate of $2.2x10^{-9}$[112], were lineage-specific and potentially undescribed. In other words, we assumed that most elements shared by large groups of mammals had already been described, and we focused instead on relatively new putative TEs. Custom scripts associated with this part of the analysis are available on github (https://github.com/davidaray/bioinfo_tools). For each iteration, we generated new consensus sequences to match the top 50 blast(*113*) hits. Bioinformatically, we accomplished this by aligning sequences with MUSCLE(*114*), trimming the alignments with trimAl with settings -gt 0.6 -cons 60(*115*), and estimating a consensus with the EMBOSS script cons with settings -plurality 3 -identity 3(*116*). We discarded files with fewer than 10 blast hits and considered consensus sequences to be "complete" if the alignments exhibited patterns of random sequence at both the 5' and 3' ends or after extending alignments to a length of 7kb or greater–whichever came first. Upon completion of this initial curation, we gathered all potential TE consensus sequences and combined them with known TEs from previous work and all known vertebrate TEs from Repbase. We then ran a cd-hit-est analysis on these sequences to identify duplicate TEs according to the 80-80-80 rule(*117*).

To confirm TE type, we compared each TE to three online databases: blastx(*118*) to confirm the presence of known ORFs in autonomous elements, RepBase(*109*) to identify known elements, and TEclass(*119*) to predict the TE type. We also used structural criteria as follows: for DNA transposons, we retained only elements with visible terminal inverted repeats. For rolling circle transposons, we required elements to have an identifiable ACTAG at one end. For putative novel short interspersed nuclear elements (SINEs), we required a repetitive tail and A and B boxes(*120*). For long terminal repeat (LTR) retrotransposons, we required recognizable hallmarks, such as TG, at their 5' ends and the reverse complement at the 3' ends.

We then combined each complete TE library with a library of known vertebrate TEs. We used Repeatmasker-4.1.0(*121*) to mask the genome assemblies with this custom library. We performed postprocessing of the outputs using a custom script, RM2Bed.py (available at https://github.com/davidaray/bioinfo_tools), which eliminates overlapping hits and converts files to .bed format.

Initial RepeatModeler output yielded 25,025 initial queries, ~101/assembly. After *de novo* curation and elimination of duplicates, the final library consisted of 7,707 consensus sequences. We have deposited these in the dfam TE database (https://dfam.org/home).

### Constraint scoring

#### *Mammalian neutral model*

We used the alignment of 241 mammals(*16*) in HAL-format (HAL Tools v2.1)(*107*) as the input to generate three different neutral models. The first was a general model, used for autosomes, and the other two were for each of the sex chromosomes (chrX and chrY). We used the resulting nucleotide substitution rate matrices to generate conservation scores. For this process, we first identified the ancestral repeats by running RepeatMasker Open-4.0. 2013-2015(*121*) on the ancestral sequence of the mammal alignment. We used sequence from the second most ancestral branch (fullTreeAnc238) instead of the most ancestral sequence (fullTreeAnc239), as interspersed repeats on fullTreeAnc238 had better reconstruction of the eutherian ancestral form than in fullTreeAnc239 (personal communication A. Smit)(*121*). We converted repeat coordinates to fullTreeAnc239 using halLiftover(*107*). Then, we filtered ancestral sequence repeat sets to exclude i) non-mammalian repeats shared with birds and reptiles; ii) repeat sequences annotated as structural RNA copies, satellites, tandem repeats, and low complexity annotations; and iii) regions where synteny was not present across the four major branches of the mammalian tree: Xenarthran, Afrotherian, Laurasiatherian and Euarchontoglires (reconstruction using halLiftover).

The input for the neutral evolution model calculation was a random set of ancestral repeat positions (100kb total bases) selected from the outputs of halLiftover. We extracted an ancestor-referenced MAF alignment (hal2maf) and used PhyloFit from Phast v1.5, with default parameters (--subst-mod REV --EM) and with the corrected tree to estimate branch lengths while fixing the tree topology(*107*, *122*). We processed the resultant alignment (mafDuplicateFilter) to filter sequences that aligned multiple times to the same region(*123*). This step avoided potential biases due to species over-alignment. For sex chromosome-specific models, we converted ancestral repeat coordinates from the HAL alignment to human hg38 coordinates (halLiftover)(*107*). We selected random sets of repeat positions from either chromosome X or Y of the human outputs of halLiftover (i.e. 100Kb as above) to calculate the neutral models specific to each sex chromosome. We constructed those models separately from the general model as they evolve at a different rate than autosomes(*124*). We later used all of these models to compute phyloP scores.

We used the same method as above to estimate primate-neutral models, with the difference being that ancestral branch reconstruction was based on the 43 primates from the alignment. For these, we used human as the reference. We evaluated synteny with six of the reconstructed branches in the primates tree (fullTreeAnc112, fullTreeAnc107, fullTreeAnc103, fullTreeAnc88, fullTreeAnc78, fullTreeAnc70). We used these primate-neutral models to compute primate-specific PhastCons scores.

*Alignment pre-processing, and alignment depth, nucleotide distribution, and branch length calculations*

We converted the HAL alignment to a species-referenced MAF-format alignment using hal2maf(*107*), and then we filtered out species duplicates in the alignment using mafDuplicateFilter (https://github.com/dentearl/mafTools) in order to avoid scoring biases due to over-alignment (*123*). We allowed one sequence per species (i.e. the best match) in each alignment block to remain, which we chose by comparing the sequence to the consensus for the block(*123*). We repeated this for multiple reference species, ultimately creating human, chimpanzee, house mouse, dog, and little brown bat reference-based alignments. For the primate-specific data, we filtered out non-primate species from the alignment using the mafSpeciesSubset command from mafTools(*123*). We collected the alignment depth and list of species aligned across the alignment using the Bio.Align package from BioPython (https://biopython.org/wiki/Multiple_Alignment_Format)(*125*). We calculated total branch lengths for each of the alignment blocks in the MAF alignment files using the tree_doctor command with the branch length (–branchlen) option from the PHAST software package(*126*).

*PhyloP and PhastCons score calculations*

<u>PhyloP constraint score calculation</u>: We used phyloP (part of the PHAST v1.5 package: https://github.com/CshlSiepelLab/phast) to calculate per-base constraint and acceleration p-values(*127*). We presented scores as $-\log_{10}$ p-values under a null hypothesis of neutral evolution, where computation involved performing a likelihood ratio test at each alignment column (--method LRT) with constraint and acceleration scores outputted (--mode CONACC, negative values indicate acceleration). We calculated phyloP scores on the human-referenced, 241-way, MAF-formatted, duplicate-filtered alignment. Scores ranged from -20.0 to 8.903 for

autosomes, -20.0 to 7.765 for chromosome X, and -20.0 to 9.280 for chromosome Y; the differences of ranges of scores between the autosomes, chromosome X, and chromosome Y are due to the differences in models being used. Scores from the dog and little brown bat-referenced 241-way MAF-formatted alignments ranged from only -20 to 8.903, as their assemblies did not include chromosome Y. Scores from the human-referenced, primates-only alignment (43-way) ranged from -20 to 1.264. Scores calculated on the primate subgroup had lower ranges than mammals, as the total branch lengths of clade-specific trees were relatively low compared to the entire mammalian tree.

PhastCons constraint score calculation: We also used PhastCons, another part of the PHAST package, which uses a phylogenetic hidden Markov model (phylo-HMM) to identify evolutionarily conserved elements(*19*, *128*). In contrast to phyloP single-base constraint, PhastCons metrics incorporate the columns of flanking bases. We calculated two outputs: a per-base constraint score and a set of conserved elements coordinates (--viterbi option). We set the model parameters to match those used to generate the PhastCons output for the UCSC 100 Species Vertebrate Multiz Alignment & Conservation (expected-length=45, target-coverage=0.3, rho= 0.31, http://genome.ucsc.edu/cgi-bin/hgTrackUi?db=hg19&g=cons100way)(*129*).

Constraint score thresholds: We computed a mammalian phyloP threshold by converting the p-values corresponding to the phyloP scores into q-values using a false discovery rate (FDR) correction(*130*) (R function *qvalue*). We classified outputs as constrained or accelerated based on the sign of the score; we considered any column with a resulting q-value ≤ 0.05 to be significantly constrained or accelerated (5% FDR, 240 mammal phyloP constraint score ≥ 2.27, 3.528% of the human genome). We took a different approach for primate constraint given the fewer species (n=43) and shorter branch lengths, as a phyloP score across primates species would be underpowered to discriminate the highly constrained bases from the background. We identified the PhastCons threshold that yielded a similar fraction of the genome under constraint as for all mammals (i.e., PhastCons base score ≥ 0.961, 3.54% of the human genome). The locations of significantly constrained bases in mammals and primates overlap, which is not surprising because the 43 primates we used are a subset of the 241 mammalian assemblies. We evaluated the mammalian phyloP threshold, the use of different scores to measure constraint in mammals and primates, and the base pair resolution of mammalian phyloP scores by comparing our results to expectations of amino acid and transcription factor motif conservation (**Figs. 2 and 3**) and by using heritability analyses of human diseases and complex traits(*17*).

Lower bound estimates of genome wide constraint: We estimated the lower bound for the fraction of sites under purifying selection across the human, chimpanzee, dog, house mouse, and little brown bat genomes (π) by comparing the empirical cumulative distribution functions (ECDF) of phyloP scores across each genome to the ECDF of ancestral repeats, following the same method detailed in(*5*) where:

$\pi = 1 - \min_s F(s)/G(s)$.

Here, F is the ECDF for all sites across the genome (a function of scores s), and G is the ECDF for ancestral repeats (i.e. sites not under any evolutionary pressure). We extracted coordinates of ancestral repeats from the UCSC repeatmasker track and filtered these to retain a set of ancestral repeats present in the four clades of the mammalian tree (Xenarthran, Afrotherian, Laurasiatherian, and Euarchontoglires)(*121*, *129*). We calculated the ECDFs of phyloP values for all positions across each genome as well as those in only each set of ancestral repeats using the ecdf function in R v.4.0.4(*131*). We excluded phyloP values below -1.5 so that the extreme left tail of the ECDFs did not heavily influence estimates of constraint.

<u>Constraint analysis</u>

*Differences between species in the proportion of genome under constraint*

We found substantial differences between the proportions of constrained sequence in different species, with little brown bat having the highest proportion. To evaluate whether this may be a result of the variation in the number of closely related species in the alignment for these species rather than a true reflection of differences in constraint, we calculated the total branch length for the set of each species and each species's nine most closely related species. The correlations between these branch lengths and either the percentage (R=-0.06; p=0.9; n=5) or the amount (R=-0.47; p=0.62; n=5) of constrained sequence were not significant.

*Singletons: isolated single-base conserved positions*

We defined a "singleton" as an isolated single base-long constraint position that is over 5bp from the closest constrained position. While a fraction of singletons are presumably truly under constraint, others might reflect errors in the alignment. To identify singletons using PhyloP scores, we collected all positions for which the phyloP score was above the phyloP 5% FDR threshold calculated using the Benjamini-Hochberg method(*132*). For each position, we computed the genomic distance to the closest neighbouring constraint position. We defined phyloP constraint positions with their closest neighbor at a distance over 5bp as isolated singletons. To identify singletons using PhastCons, we selected all single base-pair long identified conserved regions and removed those with another conserved region less than or equal to 5bp away.

*Human variation at sites under constraint*

We intersected all single nucleotide polymorphisms (SNPs) in TOPMed data freeze 8 (https://topmed.nhlbi.nih.gov/), a resource of 811 million SNPs from ~186 thousand individuals(*39*), with their phyloP scores. We compared minor allele frequencies for SNPs in constraint positions (phyloP > 2.27) to those in non-constraint positions using the Wilcoxon rank sum test in R v.4.1.1. We predicted the functional impact of all SNPs using SNPEff(*133*).

*Constraint in protein-coding sequence*

We obtained protein-coding annotations from GENCODE v.36(*134*). We selected all protein-coding transcripts and checked for consistency in the cDNA, CDS, and peptide lengths (The protein-coding length had to be 3x the peptide length, and the start codon had to be methionine.). Since most genes have many transcripts, we created a transcript hierarchy to pick one transcript per gene in this order (pickOne): MANE transcript(*135*), gnomAD canonical transcript(*136*), BUSCO transcript(*137*), protein-coding transcript, lncRNA transcript, pseudogene transcript, and other transcript. For each protein-coding transcript, we identified its structural parts: proximal promoter (500bp upstream of the TSS), TSS, 5'UTR exons, start codon, coding exons, canonical intronic splice sites, introns, stop codon, and 3'UTR exons. Some protein-coding genes do not have two UTRs, and some are intronless. We extracted phyloP scores for all positions in protein-coding genes including 5' and 3' UTRs (N=34,284,184 bp) to compare constraint between different positions within coding sequences. We defined degeneracy at each protein-coding position using International Union of Pure and Applied Chemistry (IUPAC) codes and used R v.4.1.1 to summarise mean and standard deviation phyloP scores for positions within codons, degenerate and non-degenerate positions, methionines that act as and do not act as start codons, and cysteines that form and do not form intra-peptide disulfide bridges. We obtained locations of intra-peptide disulfide bridges from the UCSC Genome Browser (https://genome.ucsc.edu/cgi-bin/hgc?hgsid=799000851_rdDAOY8aECs6ODSPVUt92mNOp7fR&c=NC_045512v2&l=14950&r=24950&o=22732&t=23137&g=unipCov2DisulfBond&i=disulf+bond)(*129*).

*Four-fold degenerate sites under constraint*

To determine whether certain transcription factors (TFs) tend to bind to constrained coding sequence [as indicated by constraint in four-fold degenerate sites (4D sites), which otherwise would not be expected to show constraint], we used bedtools intersect(*138*) to identify overlap of 4D sites under constraint (phyloP > 2.270) and ENCODE transcription factor clusters downloaded from the UCSC Genome Browser (http://hgdownload.soe.ucsc.edu/goldenPath/hg38/encRegTfbsClustered/)(*135*, *138*). We took all the TFBS with cluster score = 1000 (reflects strength of the ChIP-seq signal, which are the highest-confidence TFBS) for each TF with TFBS in the database and counted the total number of constrained and non-constrained 4D sites overlapping TFBS for each TF. We then calculated a relative difference between constrained and non-constrained 4D sites for each TF:

$$\left(\frac{DcTF}{Dc} - \frac{DnTF}{Dn}\right)\times 100$$

Where *DcTF* = number of 4D constrained sites for each TF, *Dc* = number 4D constrained sites, *DnTF* = number of 4D non-constrained sites for each TF, and *Dn* = number of 4D non-constrained sites, giving a percentage constraint excess (positive) or deficiency (negative) for each TF. We then assessed whether these TFs most commonly bind to coding sequence by intersecting all TFBS with exons using bedtools(*138*) and then, for each TF, calculating the proportion of their TFBS that overlap with exons.

*Constraint in functional elements*

We calculated constraint enrichment for several genome features (coding sequences, 5' UTRs, 3' UTRs, introns, DHS, and the five types of cCREs). We obtained coding sequences, 5' UTRs, 3' UTRs and introns from GENCODE v.36 as described above(*139*). We obtained regulatory features from ENCODE3, including DNase hypersensitive sites (DHS; 243 cell lines and tissues)(*140*) and candidate *cis*-regulatory elements (cCREs)(*38*). cCREs include canonical promoter-like signatures (PLS; ± 200 bp of GENCODE TSS, high DHS and H3K4me3 signals), proximal enhancer-like signatures (pELS; ± 2 kb of GENCODE TSS, high DHS and H3K27ac signals, low H3K4me3 signal), distal enhancer-like signatures (dELS; > 2 kb from GENCODE TSS, high DHS and H3K27ac signals), DNase-H3K4me3 elements (promoter-like biochemical signature that are not within 200bp of an annotated TSS), and CTCF-only (high DNase and CTCF and low H3K4me3 and H3K27ac).

First, we calculated constrained fractions for each feature as the number of positions with phyloP above FDR=0.05 threshold/the total number of positions. Next, we calculated constraint enrichment as the constrained fraction of the feature divided by the constrained fraction of the genome. We assessed the significance of enrichment for each feature using a $\chi^2$ test in R v.4.1.1.

### Identifying regions of high constraint

*Zoonomia ultraconserved elements (zooUCEs)*

We extracted all positions in the alignment bed files where the number of species aligned was ≥ 235 and the base was the same among all species aligned at that position. We then merged neighboring positions, creating elements of ultraconserved sequences ranging in size from 2bp to 190bp. The final set of zooUCEs contains all elements ≥ 20bp. We assessed overlap between our zooUCEs and previously defined UCEs(*41*) using bedtools intersect and the "-u" flag to report all zooUCEs that overlap the original UCEs(*138*).

*Regions of contiguous constraint (RoCCs)*

We extracted all constraint positions (phyloP ≥ 2.270) from the phyloP bed files and grouped neighboring constraint positions to create runs of constraint ≥ 2bp. We noticed that many of these runs were separated by only a single base pair below the phyloP threshold, so we merged all runs of constraint that were only separated by a single base, creating Regions of Contiguous Constraint (RoCC).

*Constraint across the genome - 100kb bins*

We divided the human genome into 100kb bins, excluding chrY, any bins < 80kb positions with phyloP values, and bins < 100kb (i.e., ends of chromosomes), for a total of 28,218 bins. We then fit a linear model of the number of constrained bases within each bin using the total number of bases with phyloP values in the bin. We included several covariates in the model to control for differences between bins that could influence estimates of constraint: the number of coding bases, the number of positions with a low number of species aligning (≤24), and a measure of mappability for each bin. For mappability, we used the "k24" UCSC Genome Browser track(*129*), the fraction of 24 x 24-mer sequences overlapping a given base (i.e., the current base ± 23 bp) that map uniquely to the genome (1 = all map uniquely, 0 = none map uniquely), and we summed the number of positions in each bin with k24 score > 0.9. We square root transformed all values. We then calculated p-values from the absolute studentized residuals for each bin using the R *pnorm* function (using absolute values meant that low p-values represent significant excess or depletion of constraint), and we computed q-values from the p-values using the R function *qvalue*. We considered any bin with q-value ≤ 0.05 to be significant. We merged adjacent significant bins using *bedtools merge* (*138*). Analysis was carried out using R v4.1.1.

*Constraint in gene deserts*

We used the set of developmental TFs identified in(*141*), which are all TFs known to be involved in the regulation of developmental processes (2,863 genes). We defined intergenic regions as all regions between ensembl genes (GRCh38.103) and calculated length and proportion of constraint (number of positions with phyloP ≥ 2.270) for each intergenic region. We labelled all intergenic regions as either "neighboring developmental TF" or "other" if they do or do not border a developmental TF, respectively. We extracted all gene deserts, defined as the longest 5% of intergenic regions, and compared constraint between gene deserts neighboring developmental TFs and gene deserts neighboring other genes using a Wilcoxon rank sum test in R v. 4.0.4. We tested for a significant difference in the proportion of constraint in ENCODE3 cCREs located within gene deserts neighboring developmental TFs versus those in other gene deserts with a Wilcoxon rank sum test in R v. 4.0.4. We assessed whether genes neighboring gene deserts were enriched for developmental TFs using Fisher's exact test ("fisher.test" function in R v.4.0.4).

Identifying Unannotated Intergenic Constraint Regions (UNICORNs)

To investigate constraint in regions lacking a functional annotation, we used the “subtract” tool in BEDTools v.2.29.2(*138*) to remove sequences with the following annotations: GENCODE v37 exons (UTRs and exons for all protein-coding genes) and promoters (TSS +/- 1kb), ENCODE3 cCREs, DHS (including TF binding sites), ChIA-PET anchors, three promoter annotation sets, and 6 enhancer annotation sets. We converted annotations to hg38 coordinates when the annotations were in other assemblies. We then identified groups of closely located constraint positions within the “unannotated” sequences: We identified any two unannotated constraint positions within 5bp of each other and retained all clusters of such positions greater than 10bp in length, giving a set of 424,179 unannotated intergenic constraint regions (UNICORNs). We generated a set of non-constraint clusters to compare with the UNICORNs by identifying all unannotated intergenic regions greater than 10bp that did not contain any constraint positions (phyloP less than 2.270 for all positions). This generated over 6 million clusters, which we randomly sampled to generate a set of 424,179 to match the number of UNICORNs.

We used the “intersect” tool in BEDTools v2.30.0(*138*) to calculate the distance of each UNICORN and non-constraint cluster to its (linearly) closest gene, cCRE, dhs, and chromatin loop anchor. We identified SNPs in each cluster from TOPMed data freeze 8(*39*) and calculated average minor allele frequency and number of SNPs per bp (0-1). We compared each of these measures between UNICORNs and matched non-constraint elements using Wilcoxon rank sum tests with a Bonferroni correction for multiple testing in R v.4.1.1 after binning elements into five bins based on size (11-20bp, 21-50bp, 51-100bp, 101-500bp, 501-1,325bp).

To determine the extent to which UNICORNs overlap open chromatin regions from neurons in specific brain regions and motor cortex cell types as well as different parts of the brain at different developmental stages, we downloaded the relevant datasets and then determined their overlap with the UNICORNs. Specifically, for open chromatin for neurons from specific brain regions, we downloaded the NeuN+ ATAC-seq(*142*) data from(*45*) and processed them as described for the neurons from primary motor cortex from this dataset in(*95*). For open chromatin from specific motor cortex neuronal cell types, we used the reproducible human motor cortex neuron open chromatin regions(*44*) described in (*94*). For open chromatin from different parts of the brain at different developmental stages, we used the file GSE149268_annotation-ocr-hg38.bed.gz downloaded from GSE149268(*43*); these open chromatin datasets came from later in development than the ENCODE fetal brain open chromatin data, and, unlike the ENCODE fetal brain open chromatin, which came from the whole embryo brain, these datasets came from specific brain regions (*38*, *143*). To compute the number of UNICORNs and nucleotides within UNICORNs covered by each of these datasets as well as by the union of these datasets, we used coverageBed from bedtools(*138*). We found that the percentage of UNICORNs covered was similar to the percentage of nucleotides within UNICORNs covered.

Constraint in repeats

We downloaded repeat annotations for the human genome from the UCSC repeatmasker(*121*) track: https://hgdownload.cse.ucsc.edu/goldenPath/hg38/database/rmsk.txt.gz. We counted the number of constraint positions in each repeat contained in this database. For each repeat class ("Simple_repeat", "Low_complexity", "rRNA", "tRNA", "DNA", "Satellite", "LINE", "RC", "srpRNA", "SINE", "Retroposon", "snRNA", "scRNA", "LTR", "RNA", and "Unknown") we calculated an estimate of excess constraint per repeat type:

$$\left(\frac{rc}{c} - \frac{rn}{n}\right)\times 100$$

where $rc$ = total constrained positions in repeat class, $c$ = total constrained positions within repeats, $rn$ = total non-constrained positions in repeat class, and $n$ = total non-constrained positions within repeats, obtaining a  percentage constraint excess (positive) or deficiency (negative) for each repeat type.

Since simple repeats showed an excess of constraint, we extracted all simple repeats from the dataset and calculated distance to the nearest protein-coding gene for each one using "bedtools closest"(*138*) and RefSeq gene annotations for GRCh38 (https://www.ncbi.nlm.nih.gov/projects/genome/guide/human/index.shtml)(*121*, *144*). We then calculated the Pearson product-moment correlation coefficient between the proportion of constraint and the distance to the nearest gene using the cor.test function in R v. 4.1.1.

Identifying genes common to subsets of Zoonomia species

Starting from all protein-coding genes from the Ensembl 99 annotation(*57*) version of the human annotation, we created a GTF file for each species by taking the human-annotated transcripts and lifting the coordinates through the Zoonomia Cactus alignment via halLiftover(*107*). We made contiguous transcripts from fragmented results returned by halLiftover using two different methods. Under both methods, we considered a transcript to be "valid" if the predicted protein sequence started with methionine, was contained on a single contig, and was within 90-110% of the length of the human reference protein. These conservative criteria may have missed gene orthologs where the start codon was in an assembly gap or where the gene had substantially diverged from the human ortholog.

For the *"zoonomia_broad" method*, we smoothed the halLiftover outputs for each protein-coding gene/species/contig combination by making a single interval from the first and last coordinates for each gene ortholog in the halLiftover output. We padded both ends by 500bp and then, for each species, extracted the genome sequence in that interval. To the longest transcript, we applied Exonerate protein2genome(*145*) to find sequences matching the human protein sequence within the specified interval. We then converted the Exonerate outputs into gtf files. If the transcript was not "valid" under the criteria described above, we repeated this process for the next-longest transcript for the gene; if there were no additional transcripts in a given species, we did not report an annotation for that gene, species combination. In the event that "valid" transcripts were found for multiple contigs, we reported only the first valid transcript found. We reported transcripts along with their identity and similarity scores (from Exonerate) on both the exon and transcript level. We also reported insertions and deletions (from Exonerate) at the exon level. For the *"zoonomia_pfenning" method*, we used the method described in (cite comparative COVID-19 paper).

Given that genes are sporadically missing from each genome due to genome quality issues, cataloging "essential" genes found in all placental mammals is not feasible. We annotated just 116 genes in all 240 Zoonomia species. When we consider only the highest-quality assembly in each order, this increases to 2,718 genes, still far fewer than the 9,226 included in the BUSCO version odb10 gene set of mammalian single-copy orthologs(*146*)

We have released our data from both methods on the following website: http://genome.ucsc.edu/cgi-bin/hgGateway?genome=Homo_sapiens&hubUrl=http://cgl.gi.ucsc.edu/data/cactus/241-mammalian-2020v2-hub/hub.txt. We have labeled each entry with the corresponding human gene name, the corresponding human transcript name, the protein ensembl ids, the gene name in the species if it was available, and the method used to generate the annotation.

Identification of *CMAH* gene loss

Determining gene loss from genome assemblies can be confounded by mis-assemblies and by assembly gaps. To avoid these confounding factors, we combined two approaches, one using short reads and one using genome assemblies, to determine whether the *CMAH* gene had been lost in mammalian genomes. We selected a reference genome with an intact, well-annotated *CMAH* gene for each of the species. For the short-read approach, we obtained publicly available Illumina paired-end reads obtained from the NCBI Short Read Archive(*147*) for each species and aligned them to their assigned reference genomes with bowtie2 using the very-sensitive-local parameter(*148*). We determined read coverage of the *CMAH* coding sequence using bedtools(*138*). For the genome assembly analysis approach, we aligned the protein sequence of the *CMAH* gene from the assigned reference genome to each genome assembly using exonerate(*145*). We evaluated results from the exonerate analysis by comparing them to the Cactus multiple genome alignment. We considered putative *CMAH* gene loss events to be cases where both the short-read alignment coverage and the exonerate alignment both indicated gene loss and where both approaches indicated loss of the same part of the gene.

Identifying olfactory receptor genes

The olfactory receptor (OR) gene family shows a high frequency of duplication events that are generally unique to each lineage. We explored the OR gene family across the Zoonomia species set, independently of alignment-based annotation. We mined all genomes for OR gene sequences using the olfactory receptor assigner (ORA)(*149*). ORA uses profile hidden Markov-models (HMMs) to identify possible OR gene sequences while also highlighting pseudogene status. We used the reference OR protein sequences from ORA to identify putative OR gene regions in each assembly with tblastn(*150*, *151*). We extracted the regions reported in the tblastn results along with an additional 500bp upstream and downstream to ensure coverage of the start and stop codon regions for which neither was reported, and we used these sequences as the input to ORA. We also used all whole-genome assemblies independently as inputs for ORA. We classified genes as “pseudogenes” if they contained in-frame stop codons or were shorter than 650bp and therefore not long enough to form the 7-transmembrane domain. We excluded any pseudogenes that were less than 200bp. We clustered identified OR genes in each species with 99% identity using cd-hit(*152*) to remove putative non-paralogous duplicates. We expected two types of duplicates to be present in the data: sequences found both with tblastn and ORA and sequences that occur more than once due to duplicate contigs in the assemblies(*71*). To ensure that all sequences truly represented OR genes, we mapped each sequence to a database of 139 mammalian annotations available on RefSeq(*144*). If the highest-scoring hit for a query sequence was not an annotated OR gene, we excluded the sequence (**Table S2**). Our reference-free approach avoids underestimating ORs in distantly related taxa and is robust to variation in genome assembly quality ( OR ~ contig N50: N=236; rho = 0.006, p-value= 0.93; OR~ scaffold N50: N=236; rho = 0.013; p-value= 0.84).

Turbinal count phenotyping

We curated species-specific numbers of olfactory turbinals from both sides of the nasal cavity(*153–173*) as well as body mass (**Table S2**)(*174*). To identify turbinal homology and compute resulting counts, we followed the nomenclature according to(*153*, *154*). Our counts include frontoturbinals, ethmoturbinals, interturbinals, and the lamina semicircularis. Though the latter is not a turbinal, it exhibits a turbinal-like morphological pattern and is covered by olfactory epithelium. We did not include nasoturbinals and the septum nasi, both of which are also partly covered by olfactory epithelium in several species(*155*, *156*, *175*), because large parts of these structures are not involved in olfaction. To date, turbinal numbers are available for 67 species of our sample. We used morphological data from only adult stages because the number and complexity of olfactory turbinals increase during postnatal growth in many species (*153*, *157*, *176*). Although the given data are available for only a small sample of species, they serve as a basis for future studies on possible associations between OR genes, phenotypes, and olfactory performance.

In our statistical analyses, we excluded certain species due to imprecise data on their turbinal morphology (e.g., *Dasypus novemcinctus* (*173*)). In case of turbinal number range, we used the average for our analyses. For instance, for the domestic dog (*Canis lupus familiaris*), whose turbinal number varies widely due to extreme breed morphology (20 up to 34, n = 27 specimens(*177*)), we used the average (27 turbinals) for breeds with varying facial length types (brachycephalic to dolichocephalic(*157*, *159*)). After filtering out species and combining measurements from the same species, we had turbinal counts for 63 mammals in Zoonomia.

Statistical analyses on olfactory structures

We associated the number of OR genes with the number of olfactory turbinals using phylolm(*78*), a modified version of phylogenetic generalized least squares(*178*) that is designed to run substantially faster, with the Zoonomia phylogenetic tree(*179*). We computed an empirical p-value by comparing our p-value to p-values from 999,999 phylogenetic permulations(*79*), permutation tests in which the general phylogenetic tree topology is preserved. We used the slightly sped-up code for continuous phylogenetic permulations described in our companion paper(*180*).

Associating gene conservation with hibernation

We investigated genomic differences between mammals that we defined as hibernators and as strict homeotherms (**Table S2**). The hibernator phenotype included any species exhibiting depressed body temperature (<18℃) that persists for >24 hours. We categorized species as homeothermic if experimental data showed no evidence of naturally occurring body temperature depression in the lab or field or if species did not recover from experimentally-induced hibernation or torpor (*181–261*). We removed n=20 species categorized as daily heterotherms (body temperature declines persist for <24 hours)(*262*) from this comparison. Additionally, we removed 45 species from downstream analysis when conflicting or no evidence of hibernation had been previously described, leaving us with 22 species defined as deep hibernators and 154 species defined as strict homeotherms.

For identifying genes that are more similar to the mammalian ancestor than they are to non-hibernators through generalized least squares (GLS) forward genomics, we partitioned the exons in the 241 mammal alignment version 1(*14*) into 100bp regions. For each element, we used PREQUEL from the PHAST package(*126*) to compare the exonic sequences of hibernating versus strict homeotherms against the sequences of a mammalian ancestor that was constructed from the multi-way alignment containing 241 mammals (*14*, *16*, *19*, *126*). We generated a list of conserved exonic elements by taking the intersection of mammalian-conserved regions and exonic boundaries, where we used the conserved regions previously identified based on Siphy-ω(*263*) and excluded those shorter than 70bp. Then, grouping the conserved exonic elements within each aligned gene, we calculated the percent of bases aligned to the ancestral reference for each species. For all species in which at least 90% of the bases aligned among all conserved elements, we calculated the percent identity (# identical / # aligned) to the ancestral reference. Note that this method does have a number of limitations–it does not account for the likely effects of a sequence difference from the common ancestor (synonymous versus nonsynonymous change or type of amino acid change if the change is nonsynonymous), and it does not account for local substitution rate (changes from the common ancestor in regions with higher substitution rates are more likely to happen by chance). We then used phylogenetic GLS with the Zoonomia phylogenetic tree to identify regions conserved in hibernators relative to the placental ancestor (FDR-adjusted GLS $p<0.05$)(*178*, *179*).

To identify genes with significant evolutionary rate shifts in hibernating mammals versus non-hibernating mammals, we used RERconverge(*90*). Such genes are putative hibernation-related genes. Genes with low relative evolutionary rates in hibernating mammals likely have increased constraint in hibernators and therefore potential functional importance, while genes with high relative evolutionary rates in hibernating mammals may be under relaxation of evolutionary constraint (implying decreased functional importance) or positive selection (implying increased functional importance). Note that this method cannot accurately infer the ancestral trait. To perform RERconverge analyses, we first obtained filtered alignments for 16,209 gene exon amino acid sequences and used them to estimate tree branch lengths(*18*, *97*). We ran RERconverge using default parameters with two species sets: the full 167 mammal species set for which hibernation phenotypes were known (21 hibernators and 146 non-hibernators). Species included were those present in the protein-coding sequence alignment whose names matched species with available hibernation phenotypes. We ran getAllResiduals and foreground2Paths with the species we used supplied for the "useSpecies" argument and "clade=all" specified to foreground2Paths so that internal branches were assigned as foreground non-hibernators based on maximum parsimony. We performed correlation analyses using the correlateWithBinaryPhenotype function with default parameters. We finally performed gene set enrichment analyses using the built-in RERconverge enrichment function fastWilcoxGMTall with Gene Ontology (GO) pathway annotations from the Molecular Signatures Database (MSigDB)(*264*, *265*). While this method enabled us to identify hundreds of genes with relative evolutionary rates associated with the evolution of hibernation, this method may have missed relevant genes because it used a human-referenced protein-coding sequence alignment, and humans are not hibernators.

Correlated traits present a challenge for forward genomics. Many hibernators are bats, who also tend to be exceptionally long-lived given their body size. We were concerned that this might be confounding our results because we obtained terms related to DNA repair, which is also involved in longevity. To confirm we had not unintentionally tested for associations to longevity or other common bat traits, we repeated our analysis in the exact same way except that we excluded 11 bat species (149 species remained) and obtained similar pathway enrichment results (Data S3).

**Supplementary Text**

Overlap with Zoonomia versus 29 mammals versus 100 vertebrates

When we overlapped the human Zoonomia constrained positions with two previous comparative genomics resources, the 29 Mammals project (*32*) and the 100-vertebrate alignment (including 58 mammals)(*266*), 73.4Mb of the mammal-constrained bases identified by Zoonomia were captured in regions of constraint in at least one of the earlier data sets, and 38.8Mb were captured in both. (see figure in line for (A) Venn diagram of overlap and (B) relative proportions found in the highest constrained regions in the Zoonomia alignment).

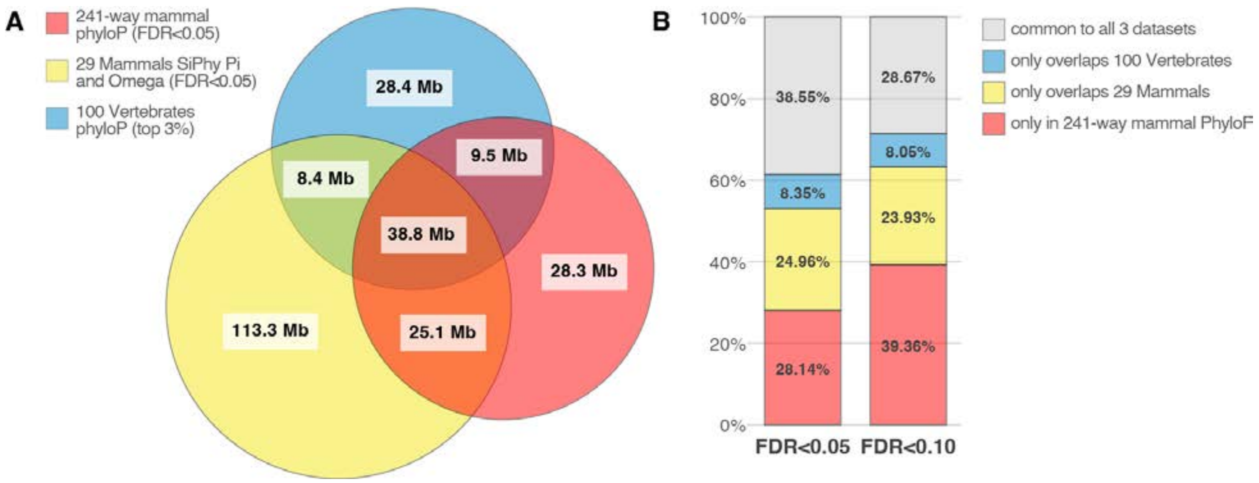

***Additional comparisons of constraint between genes and their potential regulatory elements***

We used the EpiMap putative enhancer-gene linking predictions(*267*) to link ENCODE3 proximal and distal cCREs(*38*) to their likely target genes. Briefly, EpiMap predicted enhancers in more than 800 cell types (including ENCODE datasets) and linked these putative enhancers to their target genes using correlation between enhancer presence and gene expression across these cell types. Using EpiMap enhancer-gene links in all cell types, we were able to link 33% and 53% of proximal and distal cCREs, respectively, to at least one gene.

We compared these region groupings to each other and observed some expected trends that have not previously been demonstrated at this scale. There is a strong positive correlation (Spearman rho=0.52) between conservation of protein-coding and proximal promoter sequences. This result is consistent with the theory that, if the function of a gene in mammals requires high conservation of protein sequence, then the non-coding sequence in the proximal promoter that regulates its expression also tends to be constrained(*268*, *269*). We also observed consistent correlation of protein sequence constraint with constraint in corresponding 5'UTRs and 3'UTRs (Spearman's rho = 0.54 and 0.45, respectively), and, to a lesser extent, with constraint in introns (Spearman's rho = 0.30).

To further investigate the relationship between protein-coding and regulatory conservation, we additionally evaluated the relationship between constraint in ENCODE3 cCREs(*38*) and constraint in the genes that cCREs may regulate. Investigating enhancers is challenging because cCREs do not necessarily regulate the closest genes(*270*) and because ENCODE3 did not provide any cCRE-gene linking predictions. We observed non-negligible correlation between distal cCRE constraint scores and constraint in associated CDSs and promoters (Spearman's rho = 0.20 and 0.23, respectively).

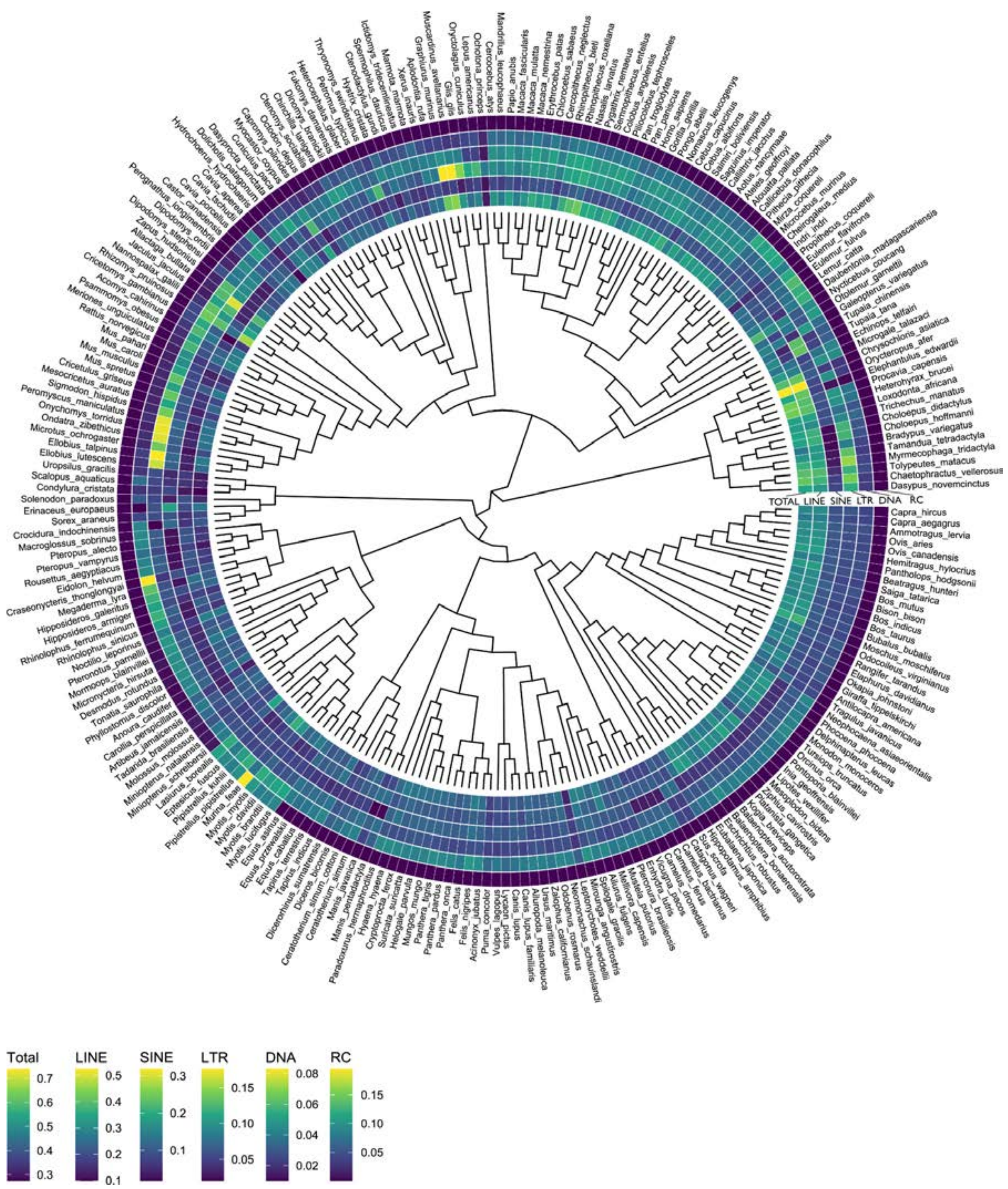

**Fig. S1. Proportion of genomic content attributed to transposable element accumulation within a phylogenetic context.** The rings, from inner to outer, depict the transposable element accumulation data as proportions of the total genome assembly: all transposable element content, LINEs, SINEs, LTRs, DNA transposons, rolling-circle transposons. Cladogram adapted from (*271*).

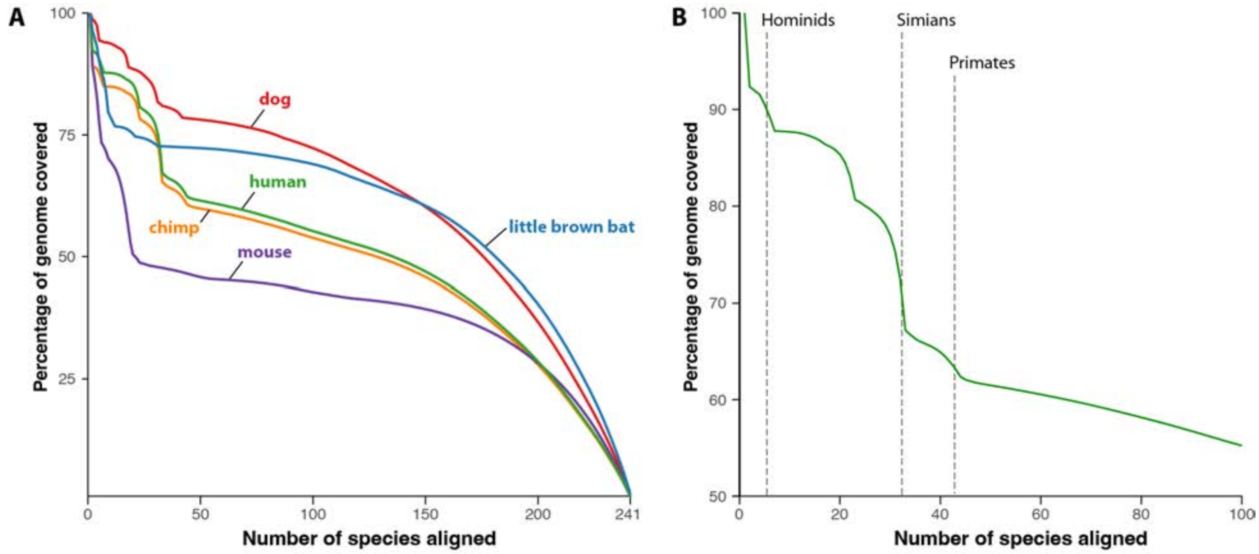

**Fig. S2. Genome alignment varies among Zoonomia species. (A)** When the Cactus alignment is referenced on five different species, the depth of the alignment (number of species aligned) varies widely. Only a small percentage of the genome aligns in most species, with rapid drops correlating with longer branches of the phylogeny. **(B)** For example, in the human alignment, most of the genome is aligned in 5 other hominids but this drops as more species, and more distantly related species, are included.

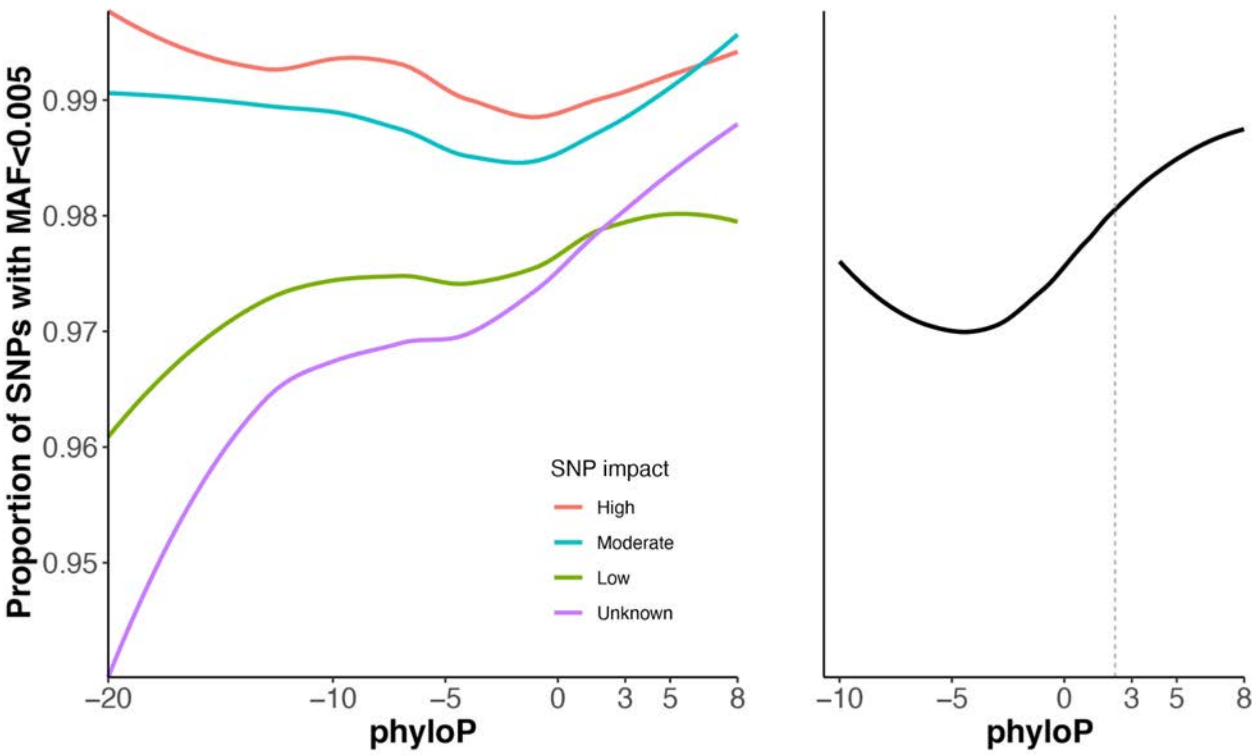

**Fig. S3. phyloP scores versus percentage of single-nucleotide variants (SNPs) with minor allele frequency (MAF) < 0.005. (A)** phyloP score versus percentage of SNPs with MAF < 0.005 for SNPs with different predicted impact levels by SNPEff. **(B)** phyloP score versus percentage of SNPs with MAF < 0.005 for all SNPs. SNPs are from TOPMed data freeze 8(*39*).

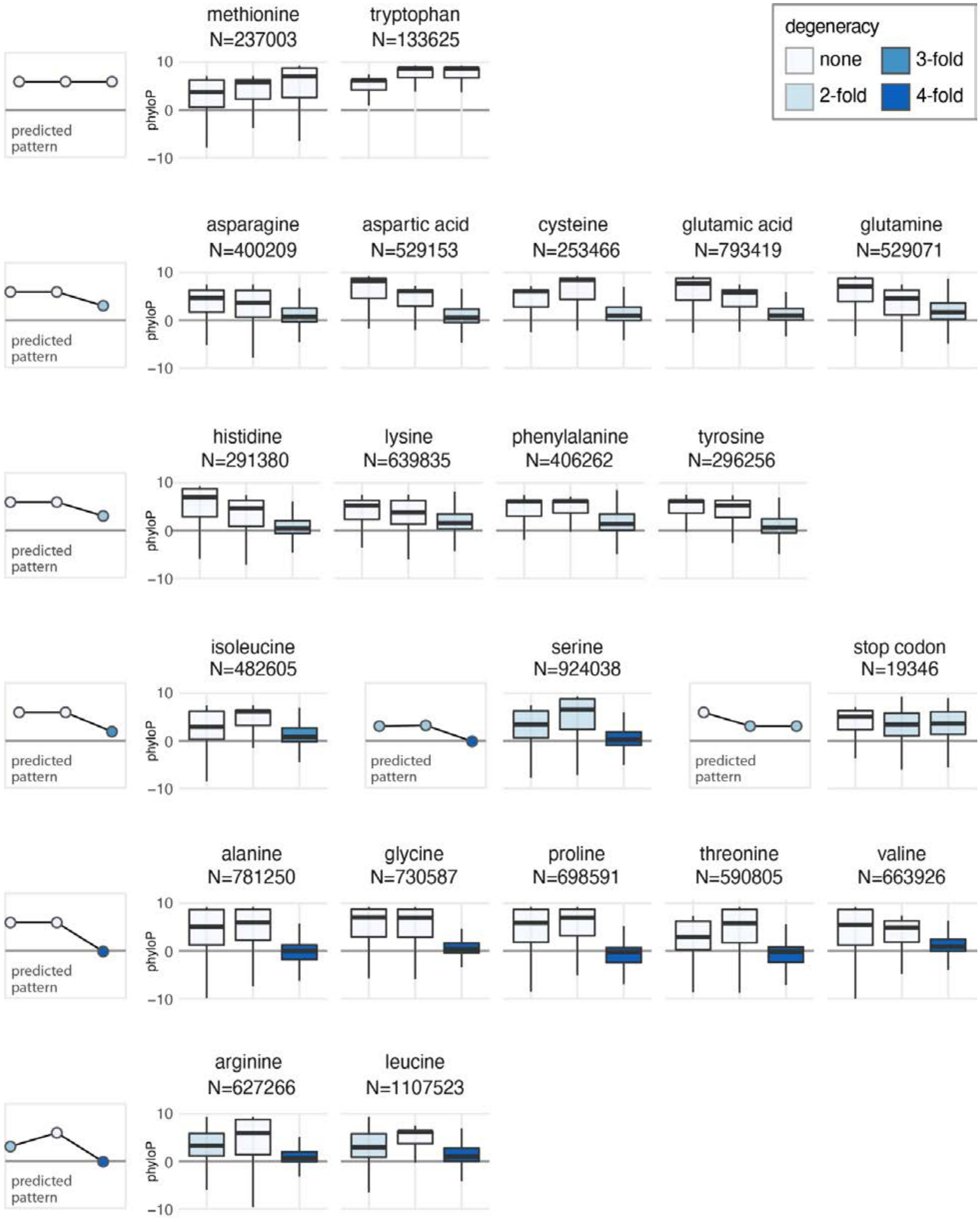

**Fig. S4. Degeneracy and constraint in codon positions.** Boxplots showing the distribution of phyloP at each position within a codon for all 20 amino acids plus stop codon. Predicted patterns of constraint are based on the degeneracy for each amino acid (ranging from 0 (four-fold degenerate) to 4 (non-degenerate). They appear similar to the observed patterns of constraint.

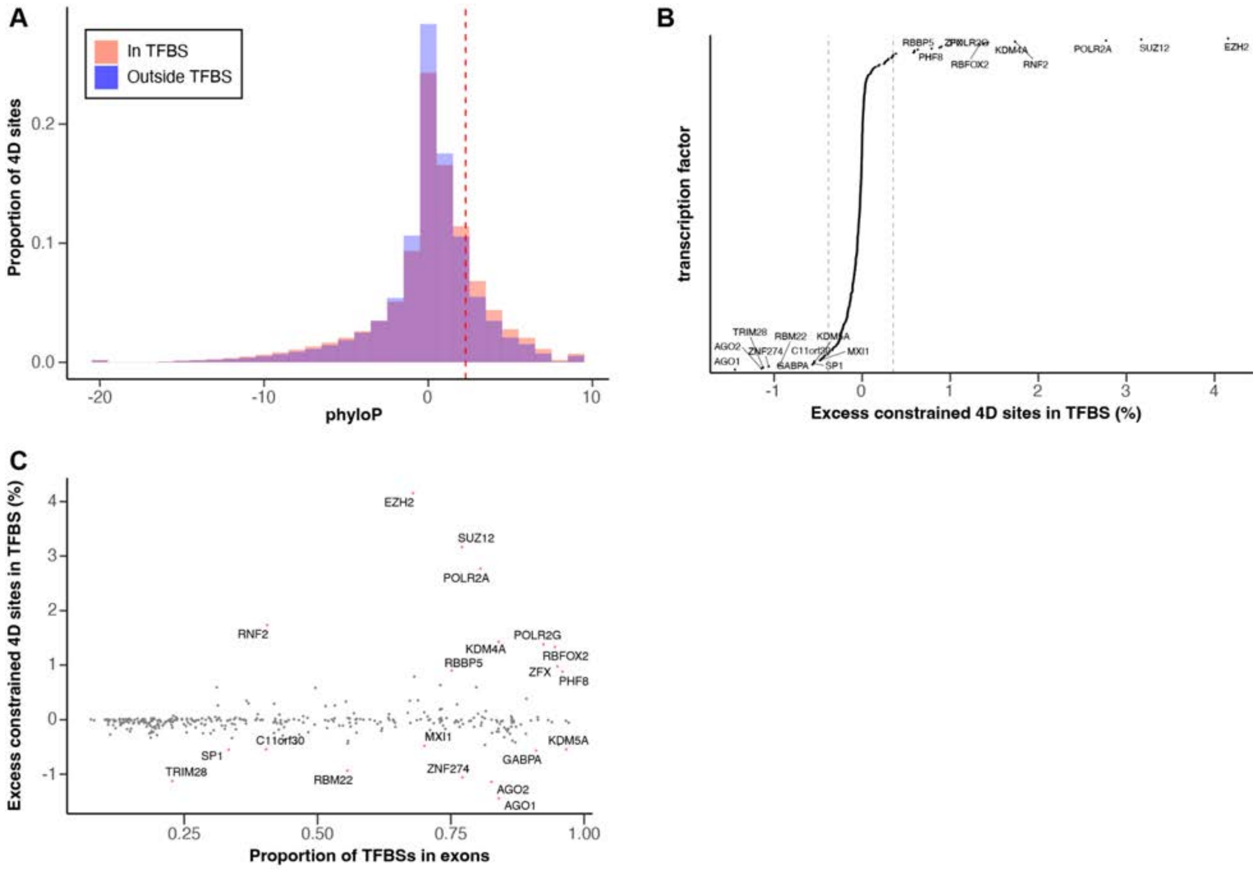

**Fig. S5. Constraint and transcription factor binding at four-fold degenerate sites. (A)** phyloP scores at four-fold degenerate sites inside versus outside of TFBS. **(B)** Level of excess constraint at four-fold degenerate sites in TFBS for each transcription factor. Dashed vertical lines represent 5th and 95th percentiles. **(C)** Proportion of TFBS in exons versus excess constraint at four-fold degenerate sites for transcription factors. In **(B)** and **(C)**, labelled transcription factors have notably high or low levels of excess constraint at four-fold degenerate sites.

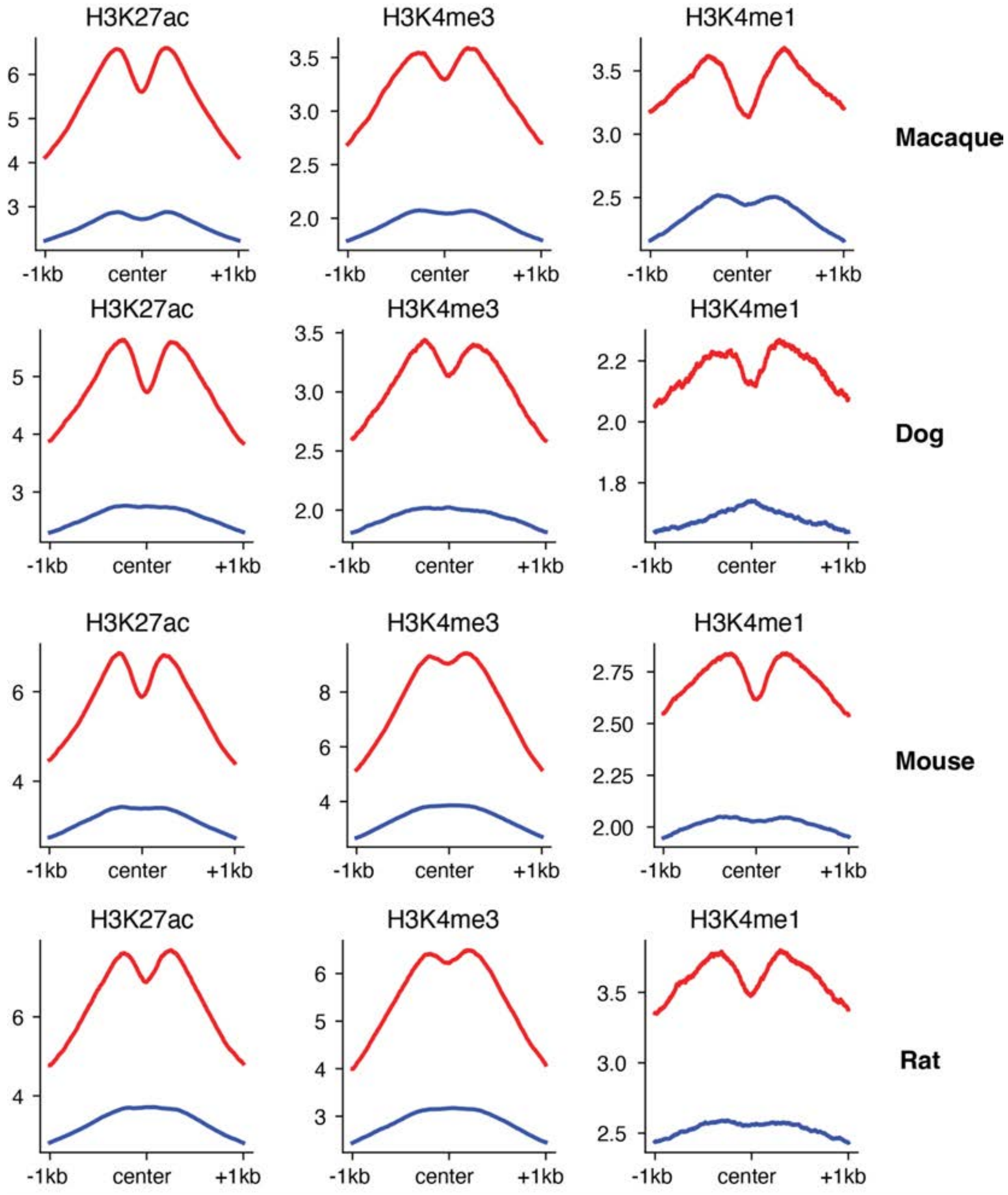

**Fig. S6. Constraint in TFBS across species.** Active human histone marks H3K4me3, H3K27ac, and H3K4me1 are enriched at constrained TFBS in other species (red) but not at unconstrained TFBS (blue). Y-axis shows enrichment.

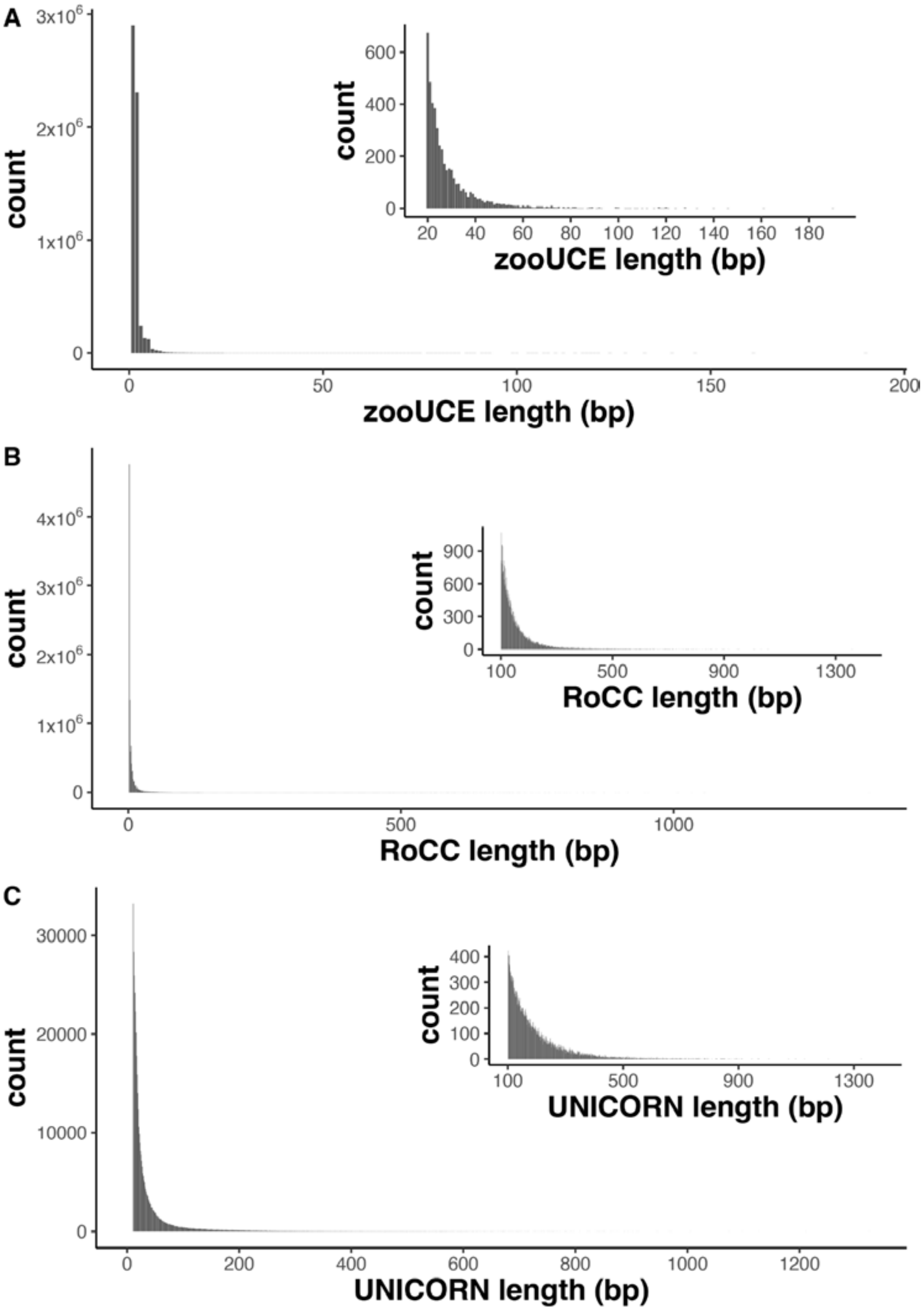

**Fig. S7. Histograms of lengths of conserved regions for different definitions of conservation.** **(A)** Histogram of lengths of Zoonomia ultraconserved elements (zooUCEs). **(B)** Histogram of lengths of regions of contiguous conservation (RoCCs). **(C)** Histogram of lengths of UNICORNs. Insets show distributions for shorter length elements.

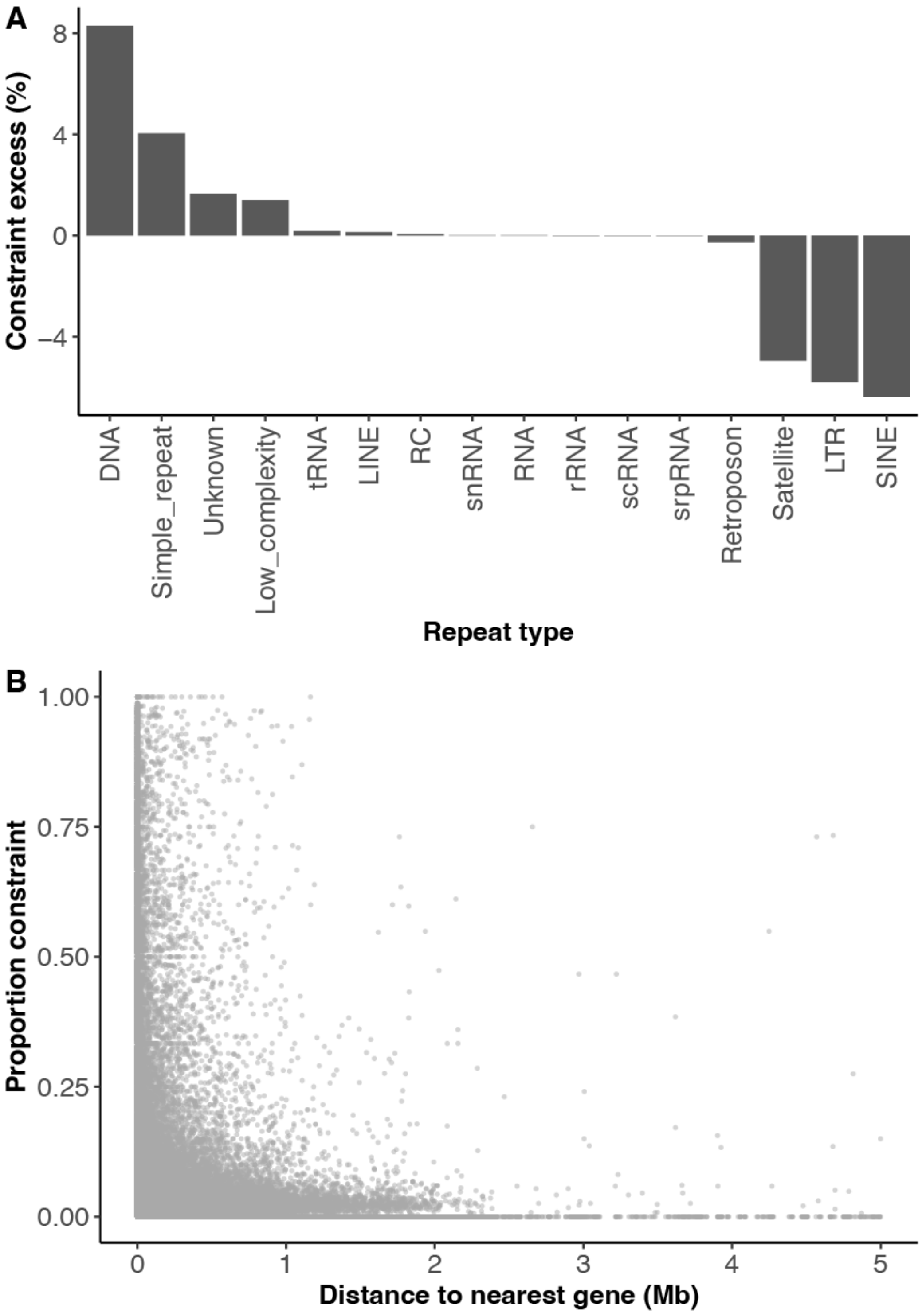

**Fig. S8. Constraint of repeats. (A)** Percentage of excess or depletion of constraint in each human repeat class. **(B)** Distance to nearest protein-coding transcription start site versus proportion of positions under constraint (phyloP > 2.27) for all human simple repeats.

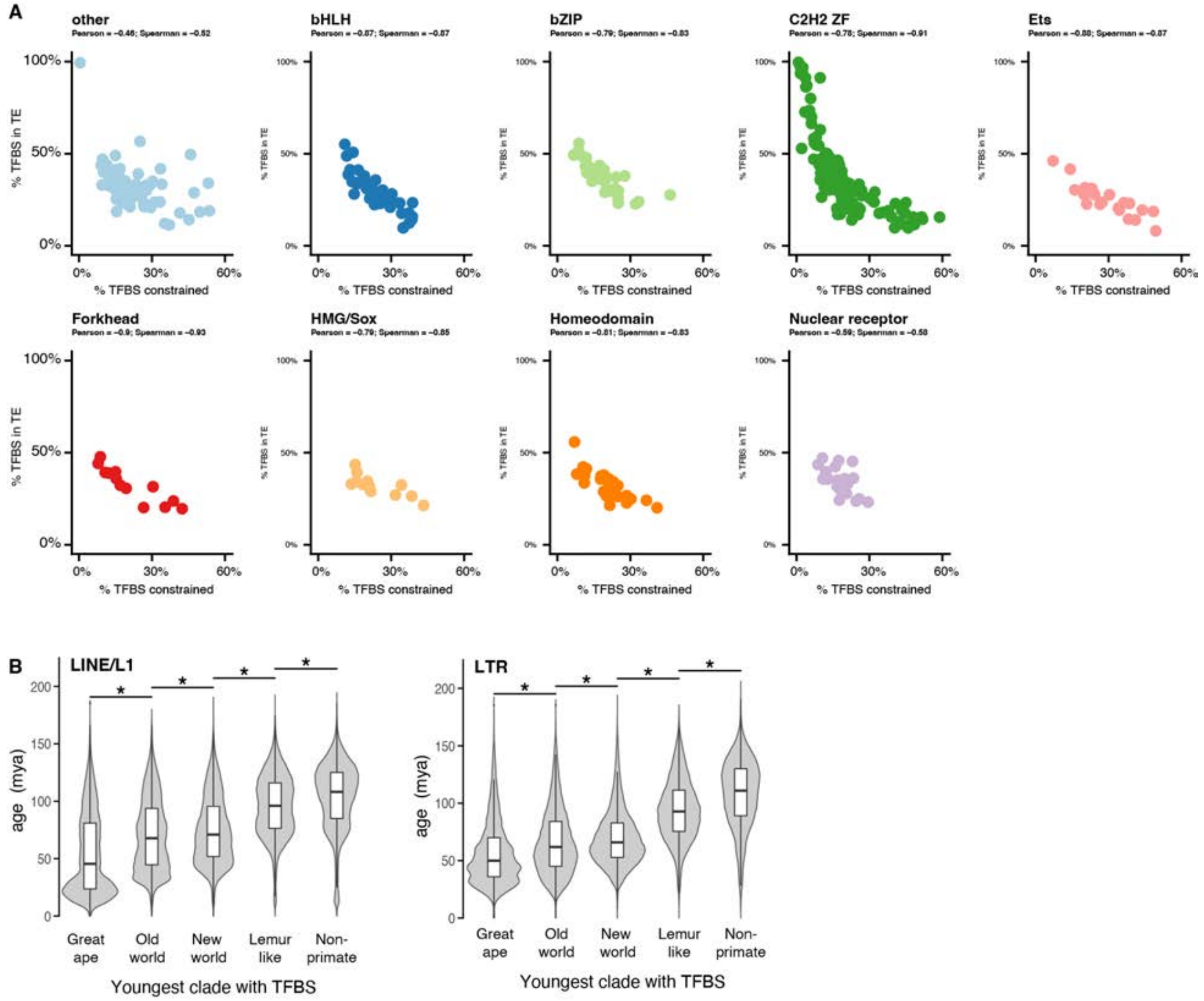

**Fig. S9. Transcription factor binding sites (TFBS) in transposable elements. (A)** Percent of TFBS found in a transposable element versus the percent found to be evolutionary constrained for 368 transcription factors, separated by family. **(B)** TFBS found share only in more closely related species tend to be found on younger transposable elements, and vice versa.

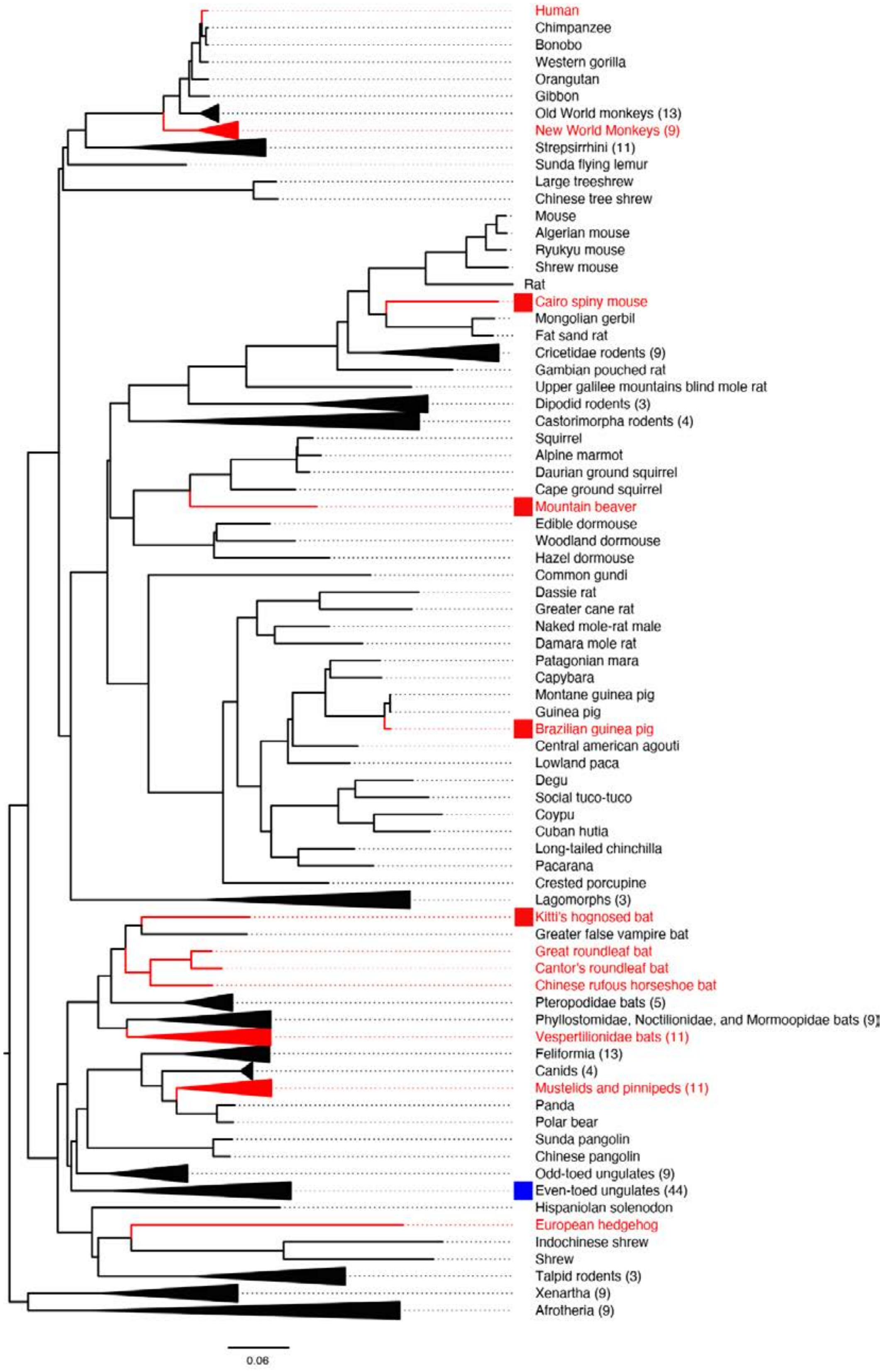

**Fig. S10. Phylogenetic tree with CMAH gene loss annotated.** Red lineages indicate CMAH loss.

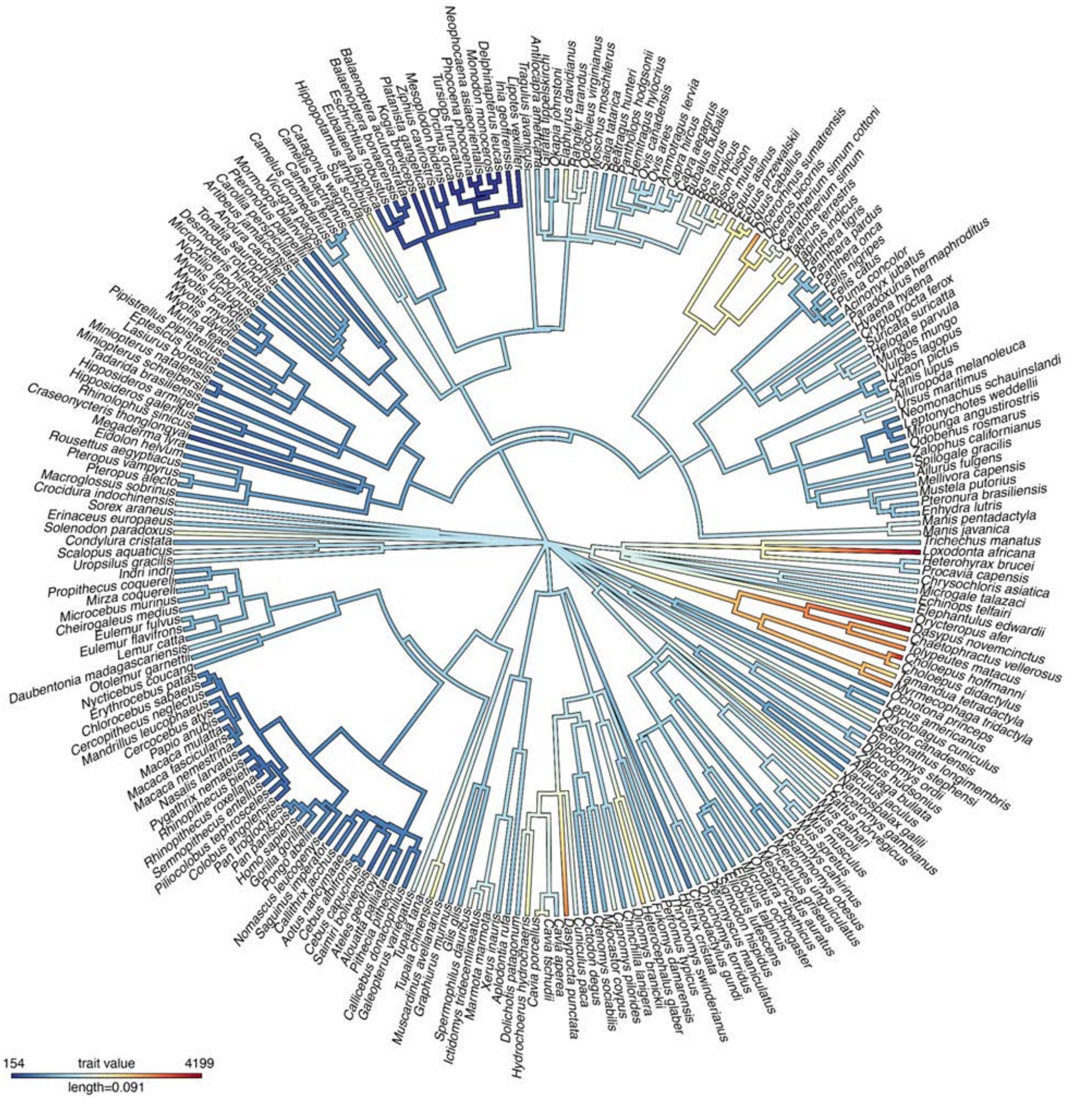

**Fig. S11. Phylogenetic tree annotated with the number of olfactory receptor genes in each species** (trait is number of olfactory receptors, both functional and non-functional).

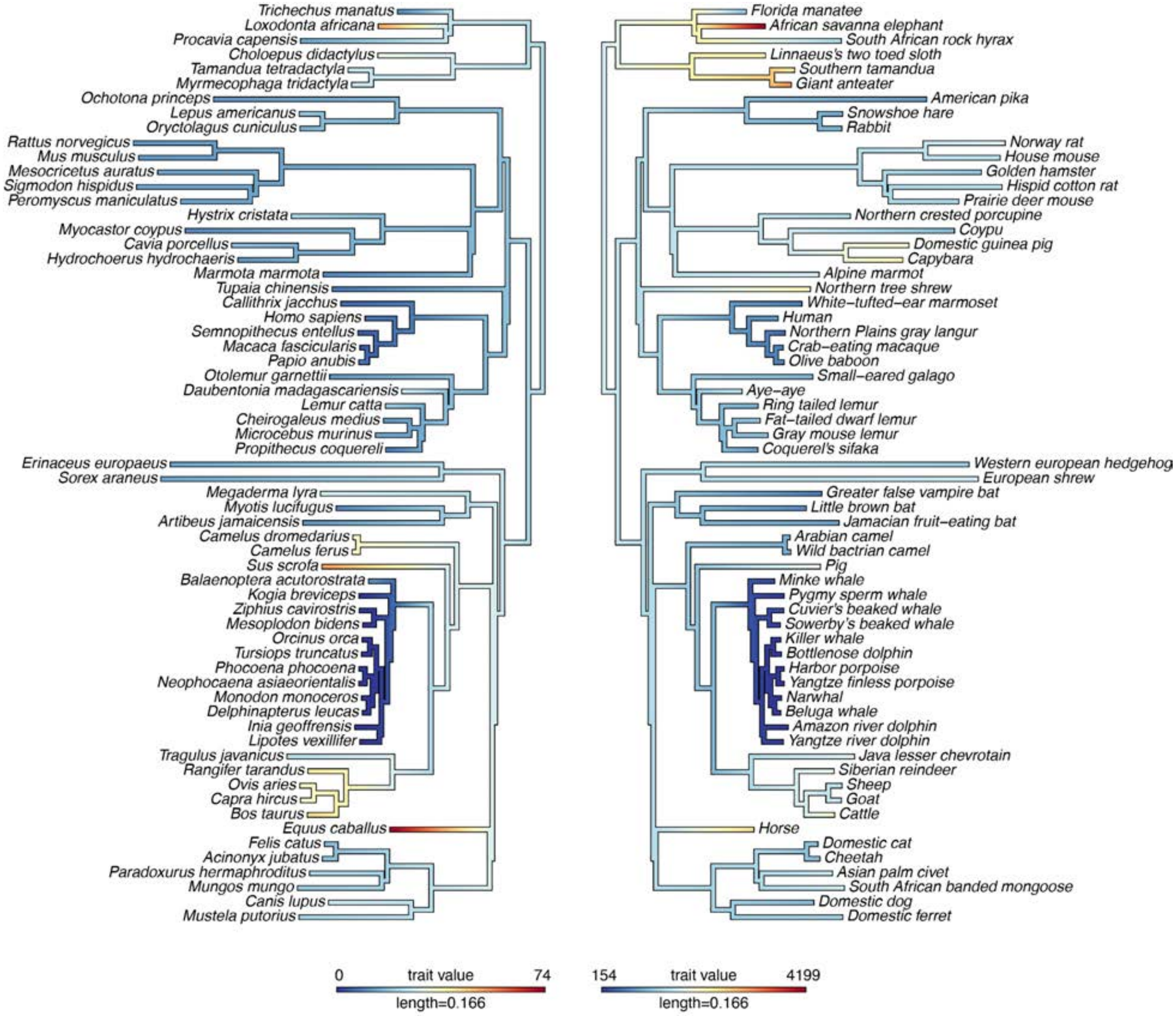

**Fig. S12. Phylogenetic trees annotated with olfactory receptor gene counts (left) and number of olfactory turbinals (right).** Species included are the subset of species with both olfactory receptor gene counts and number of olfactory turbinal annotations. Species' binomial names are listed on the left, and species' common names are listed on the right.

**Table S1.**

Genome-wide constraint and proportions of the genome under constraint at different false discovery rates for human, chimpanzee, mouse, dog, and little brown bat.

| species | Genome assembly | % genome scored | | Total % constrained | < 1% FDR | < 3% FDR | < 5% FDR* | < 10% FDR | < 20% FDR |
|---|---|---|---|---|---|---|---|---|---|
| Human (*Homo sapiens*) | GRCh38.p12 (3.09Gb) | 92.4% | phyloP cutoff | n/a | 3.27 | 2.59 | 2.27 | 1.78 | 1.25 |
| | | | % genome | 13.1% | 1.87% | 2.67% | 3.26% | 4.69% | 7.56% |
| | | | bases (Mb) | 405 | 58 | 82 | 101 | 145 | 233 |
| Chimpanzee (*Pan troglodytes*) | panTro6 (3.10Gb) | 89.1% | phyloP cutoff | n/a | 3.26 | 2.59 | 2.27 | 1.78 | 1.25 |
| | | | % genome | 13.8% | 1.8% | 2.6% | 3.2% | 4.6% | 7.4% |
| | | | bases (Mb) | 428 | 57 | 81 | 99 | 142 | 229 |
| Mouse (*Mus musculus*) | GRCm38.p6 (2.73Gb) | 89.1% | phyloP cutoff | n/a | 3.63 | 3.02 | 2.70 | 2.27 | 1.79 |
| | | | % genome | 11.1% | 1.7% | 2.1% | 2.6% | 3.2% | 4.4% |
| | | | bases (Mb) | 303 | 47 | 58 | 71 | 88 | 120 |
| Dog (*Canis lupus familiaris*) | CanFam3.1 (2.33Gb) | 99.0% | phyloP cutoff | n/a | 3.52 | 2.88 | 2.56 | 2.10 | 1.59 |
| | | | % genome | 11.4% | 2.11% | 2.84% | 3.43% | 4.69% | 7.02% |
| | | | bases (Mb) | 265 | 49.2 | 66.0 | 79.8 | 109.2 | 163.4 |
| Little brown bat (*Myotis lucifugus*) | Myoluc2.0 (2.03Gb) | 96.5% | phyloP cutoff | n/a | 3.48 | 2.85 | 2.54 | 2.09 | 1.61 |
| | | | % genome | 19.8% | 2.45% | 3.27% | 3.85% | 5.16% | 7.38% |
| | | | bases (Mb) | 403 | 50 | 67 | 78 | 105 | 150 |

* a "constrained base" is defined as one with a phyloP score above the 5% FDR cutoff.

***Table S2.***

Species' names, species' taxonomy, analyses in which species were included, and species' phenotype information. "NA" means data was not available or not used in analysis. Annotations in parentheses indicate that neither the phenotype annotation information nor the assembly in the Cactus alignment had an indication of subspecies, but the TOGA alignments have two subspecies. The "Brain size residual" is the residual of the brain size when regressing out body size.

**Table S3.**
Five regions of the human genome are assigned to mammalian multi-species homologous synteny blocks significantly larger than expected by chance. In total, 44% (1.3 Gb) of the human genome was assigned to multi-species homologous synteny blocks (msHSBs) at least 1 Mb long and present in all mammalian species. Five msHSBs are longer than expected by chance ($P < 0.05$; >10 Mb) (in hg38 coordinates).

| chr | start | end | length | p-value |
|---|---|---|---|---|
| chr2 | 160984329 | 183095659 | 22111330 | 1.09E-06 |
| chr8 | 63166966 | 75150359 | 11983393 | 0.0103 |
| chr8 | 97763558 | 109715243 | 11951685 | 0.0106 |
| chr9 | 8064120 | 19126337 | 11062217 | 0.0235 |
| chr5 | 86356760 | 96585715 | 10228955 | 0.0492 |

**Table S4.**
Putative horizontally transferred transposable elements involving non-chiropterans. Outside of bats, we found 11 instances of potential TE introduction through horizontal transfer (transfer from one species to another in the absence of reproduction). In contrast, in bats, we identified 222 putative horizontal transfers, including Tc-Mariner, hAT(8), and piggyBac elements.

| **Element** | **TE Family** | **Autonomy** | **Library Consensus Length (bp)** | **Species Involved** | **Full-Length Hits** |
|---|---|---|---|---|---|
| HysCri-1.134 | hAT | Non-autonomous | 192 | *Heterocephalus glaber* | 303 |
| | | | | *Hystrix cristata* | 257 |
| nhAT2_ML | hAT | Non-autonomous | 204 | *Cheirogaleus medius* | 91 |
| | | | | *Nycticebus coucang* | 439 |
| NycCou-1.114 | hAT | Non-autonomous | 215 | *Nycticebus coucang* | 123 |
| PMER1 | hAT | Non-autonomous | 90 | *Otolemur garnettii* | 1857 |
| npiggy1_Mm | PiggyBac | Non-autonomous | 240 | *Microcebus murinus* | 890 |
| | | | | *Mirza coquereli* | 518 |
| ScaAqu-1.134 | PiggyBac | Non-autonomous | 520 | *Scalopus aquaticus* | 453 |
| ScaAqu-1.148 | PiggyBac | Non-autonomous | 240 | *Scalopus aquaticus* | 143 |
| ScaAqu-1.172 | PiggyBac | Non-autonomous | 517 | *Scalopus aquaticus* | 225 |
| HipAmp-1.103 | Tc-Mariner | Non-autonomous | 966 | *Hippopotamus amphibius* | 368 |
| OdoVir-5.847 | Tc-Mariner | Autonomous | 1283 | *Balaenoptera acutorostrata* | 985 |
| | | | | *Balaenoptera bonaerensis* | 398 |
| | | | | *Delphinapterus leucas* | 922 |
| | | | | *Eschrichtius robustus* | 524 |
| | | | | *Eubalaena japonica* | 511 |
| | | | | *Hippopotamus amphibius* | 332 |
| | | | | *Inia geoffrensis* | 483 |
| | | | | *Kogia breviceps* | 477 |
| | | | | *Lipotes vexillifer* | 728 |
| | | | | *Mesoplodon bidens* | 490 |
| | | | | *Monodon monoceros* | 516 |
| | | | | *Neophocaena asiaeorientalis* | 820 |
| | | | | *Orcinus orca* | 875 |
| | | | | *Phocoena phocoena* | 521 |
| | | | | *Platanista gangetica* | 484 |

| | | | | | |
|---|---|---|---|---|---|
| | | | | *Pontoporia blainvillei* | 277 |
| | | | | *Tursiops truncatus* | 688 |
| | | | | *Ziphius cavirostris* | 464 |
| ProCoq-1.279 | Tc-Mariner | Non-autonomous | 131 | *Cheirogaleus medius* | 170 |
| | | | | *Daubentonia madagascariensis* | 413 |
| | | | | *Eulemur flavifrons* | 168 |
| | | | | *Eulemur fulvus* | 192 |
| | | | | *Indri indri* | 220 |
| | | | | *Lemur catta* | 168 |
| | | | | *Microcebus murinus* | 117 |
| | | | | *Mirza coquereli* | 127 |
| | | | | *Propithecus coquereli* | 164 |

**Table S5.**
Regions (≥100 Kb) in the human genome with significantly high (positive standardized residual) or low (negative standardized residual, only one window) constraint (q<0.05 from a linear model) and genes within those bin.

| Chr | Start | End | Length | q-value | Standardized residuals | Fraction conserved | Fraction CDS | Gene |
|---|---|---|---|---|---|---|---|---|
| chr2 | 144400000 | 144600000 | 200000 | 0.000022 | 6.15,5.94 | 0.2318 | 0.0081 | ZEB2 |
| chr7 | 27100000 | 27200000 | 100000 | 0.000025 | 6.02 | 0.3333 | 0.0974 | HOXA |
| chr2 | 60400000 | 60600000 | 200000 | 0.000066 | 4.19,5.74 | 0.1916 | 0.0156 | BCL11A |
| chr12 | 53900000 | 54100000 | 200000 | 0.00014 | 4.31,5.58 | 0.2150 | 0.0384 | HOXC |
| chr8 | 76600000 | 76700000 | 100000 | 0.000159 | 5.52 | 0.1452 | 0.0000 | ZFHX4 |
| chr2 | 66400000 | 66700000 | 300000 | 0.000387 | 4.26,5.33,4.4 | 0.1447 | 0.0055 | MEIS1 |
| chr19 | 8900000 | 9000000 | 100000 | 0.000395 | -5.31 | 0.0077 | 0.3895 | MUC16 |
| chr1 | 10700000 | 10800000 | 100000 | 0.000442 | 5.26 | 0.1720 | 0.0000 | CASZ1 |
| chr2 | 176100000 | 176200000 | 100000 | 0.000754 | 5.15 | 0.2626 | 0.0763 | HOXD |
| chr2 | 58900000 | 59000000 | 100000 | 0.000992 | 5.08 | 0.1539 | 0.0000 | LINC01122 |
| chr10 | 129800000 | 130000000 | 200000 | 0.001091 | 4.79,5.03 | 0.1672 | 0.0094 | EBF3 |
| chr15 | 36900000 | 37100000 | 200000 | 0.001091 | 4.35,5.04 | 0.1693 | 0.0055 | MEIS2 |
| chr18 | 25100000 | 25200000 | 100000 | 0.001778 | 4.92 | 0.1736 | 0.0011 | ZNF521 |
| chr1 | 27500000 | 27600000 | 100000 | 0.00216 | 4.84 | 0.2055 | 0.0481 | AHDC1 |
| chr3 | 114400000 | 114500000 | 100000 | 0.00216 | 4.86 | 0.1675 | 0.0000 | ZBTB20 |
| chr16 | 73000000 | 73100000 | 100000 | 0.00216 | 4.85 | 0.1498 | 0.0000 | ZFHX3 |
| chr1 | 87300000 | 87400000 | 100000 | 0.00278 | 4.77 | 0.1804 | 0.0049 | LMO4 |
| chr10 | 101400000 | 101500000 | 100000 | 0.003673 | 4.7 | 0.1271 | 0.0027 | BTRC |
| chr17 | 48500000 | 48600000 | 100000 | 0.005084 | 4.62 | 0.2217 | 0.0596 | HOXB |
| chr3 | 169200000 | 169500000 | 300000 | 0.006499 | 4.56,4.03,4.29 | 0.1317 | 0.0011 | MECOM |
| chr2 | 143300000 | 143400000 | 100000 | 0.008396 | 4.5 | 0.1404 | 0.0000 | ARHGAP15 |
| chr7 | 114600000 | 114700000 | 100000 | 0.010962 | 4.42 | 0.1864 | 0.0198 | FOXP2 |
| chr9 | 106800000 | 106900000 | 100000 | 0.010962 | 4.42 | 0.1160 | 0.0000 | ZNF462 |
| chr10 | 113100000 | 113200000 | 100000 | 0.010962 | 4.42 | 0.1735 | 0.0151 | TCF7L2 |
| chr15 | 96300000 | 96400000 | 100000 | 0.010969 | 4.4 | 0.1683 | 0.0128 | NR2F2 |
| chr2 | 143600000 | 143700000 | 100000 | 0.012204 | 4.36 | 0.1347 | 0.0013 | ARHGAP15 |
| chr15 | 36600000 | 36700000 | 100000 | 0.012204 | 4.36 | 0.1353 | 0.0047 | C15orf41 |
| chr10 | 75700000 | 75800000 | 100000 | 0.012856 | 4.33 | 0.1342 | 0.0005 | LRMDA |
| chr2 | 172000000 | 172100000 | 100000 | 0.013449 | 4.32 | 0.1600 | 0.0173 | DLX1-2 |
| chr9 | 125700000 | 125900000 | 200000 | 0.013763 | 4.3,4.21 | 0.1333 | 0.0015 | MAPKAP1 |
| chr5 | 158900000 | 159100000 | 200000 | 0.013874 | 4.29,4.27 | 0.1500 | 0.0027 | EBF1 |
| chr14 | 33600000 | 33700000 | 100000 | 0.014583 | 4.25 | 0.1501 | 0.0030 | NPAS3 |
| chr15 | 84400000 | 84500000 | 100000 | 0.014583 | 4.25 | 0.0979 | 0.0000 | |

| | | | | | | | | |
|---|---|---|---|---|---|---|---|---|
| chr17 | 20700000 | 20800000 | 100000 | 0.014583 | 4.25 | 0.0911 | 0.0000 | |
| chr2 | 163700000 | 163800000 | 100000 | 0.016727 | 4.19 | 0.1316 | 0.0002 | FIGN |
| chr5 | 139600000 | 139700000 | 100000 | 0.016727 | 4.2 | 0.1468 | 0.0138 | CXXC5 |
| chr10 | 21500000 | 21600000 | 100000 | 0.016727 | 4.19 | 0.1409 | 0.0366 | MLLT10 |
| chr10 | 61900000 | 62100000 | 200000 | 0.017361 | 4.18,3.96 | 0.1551 | 0.0179 | ARID5B |
| chr5 | 88600000 | 88700000 | 100000 | 0.018096 | 4.16 | 0.1112 | 0.0000 | LINC00461 |
| chr3 | 71000000 | 71100000 | 100000 | 0.018405 | 4.15 | 0.1510 | 0.0080 | FOXP1 |
| chr5 | 141800000 | 141900000 | 100000 | 0.019259 | 4.14 | 0.1511 | 0.0391 | PCDH1 |
| chr1 | 87500000 | 87600000 | 100000 | 0.021706 | 4.11 | 0.1100 | 0.0000 | |
| chr3 | 62400000 | 62500000 | 100000 | 0.023289 | 4.09 | 0.1596 | 0.0129 | CADPS |
| chr2 | 178500000 | 178600000 | 100000 | 0.02534 | 4.06 | 0.3759 | 0.5223 | TTN |
| chr1 | 62900000 | 63000000 | 100000 | 0.025537 | 4.05 | 0.1280 | 0.0000 | LINC01739 |
| chr9 | 123700000 | 123800000 | 100000 | 0.025537 | 4.05 | 0.1322 | 0.0021 | DENND1A |
| chr2 | 63000000 | 63100000 | 100000 | 0.02831 | 4.02 | 0.1339 | 0.0155 | OTX1 |
| chr1 | 90800000 | 90900000 | 100000 | 0.031255 | 3.99 | 0.0916 | 0.0000 | LINC02609 |
| chr10 | 100600000 | 100800000 | 200000 | 0.03856 | 3.93,3.85 | 0.1252 | 0.0037 | PAX2 |
| chr1 | 44200000 | 44300000 | 100000 | 0.047713 | 3.88 | 0.1514 | 0.0202 | DMAP1 |
| chr2 | 59900000 | 60000000 | 100000 | 0.047713 | 3.87 | 0.1259 | 0.0000 | |
| chrX | 74100000 | 74200000 | 100000 | 0.047713 | 3.87 | 0.0532 | 0.0000 | AL359740.1 |
| chr3 | 181700000 | 181800000 | 100000 | 0.047781 | 3.86 | 0.1306 | 0.0095 | SOX2 |
| chr13 | 72500000 | 72600000 | 100000 | 0.047781 | 3.87 | 0.1171 | 0.0000 | |

**Table S6.**
Annotation sources used for identifying genomic regions outside of annotations when defining UNICORNs.

| **Annotation type** | **Name** | **Date accessed** | **Details** | **Link** |
|---|---|---|---|---|
| **Coding regions** | GENCODE v37 | 05.21.2021 | UTRs and exons for all protein-coding genes | https://www.gencodegenes.org/human/release_37.html |
| | | | Promoters (TSS +/- 1kb) | Manually calculated from each TSS |
| **Regulatory features** | ENCODE3 cCREs | 06.03.2021 | Candidate *cis*-regulatory elements, including promoter-like signatures (PLS), proximal enhancer-like signatures (pELS) and distal enhancer-like signatures (dELS) | https://screen.encodeproject.org/ |
| | ENCODE3 DHS | 06.03.2021 | DNase hypersensitive sites in 243 cell lines | https://doi.org/10.1038/s41586-020-2528-x |
| | ENCODE3 ChIA-PET anchors | 06.03.2021 | Chromatin loop anchors identified in 24 cell types | https://doi.org/10.1038/s41586-020-2151-x |
| | UCSC Promoters | 08.20.2021 | Experimentally validated promoters generated by the Eukaryotic Promoter Database. | https://genome.ucsc.edu/cgi-bin/hgTrackUi?db=mm10&c=chrX&g=epdNew |
| | Human_promoter_Villar | 08.20.2021 | Promoters active in human liver | https://doi.org/10.1016/j.cell.2015.01.006 |
| | Enhancer_Andersson | 08.20.2021 | Atlas of active enhancers across human cell types and tissues | https://doi.org/10.1038/nature12787 |
| | Enhancer_Hoffman | 08.20.2021 | Enhancers identified from ENCODE data | https://doi.org/10.1093/nar/gks1284 |
| | Human_Enhancer_Villar | 08.20.2021 | Enhancers active in human liver | https://doi.org/10.1016/j.cell.2015.01.006 |
| | SuperEnhancer_Hnisz | 08.20.2021 | Catalog of super-enhancers in 86 human cell and tissue types | https://doi.org/10.1016/j.cell.2013.09.053 |

**Table S7**
Genome assembly quality and predicted gene statistics.

| **Pearson correlation** | | | | | | |
|---|---|---|---|---|---|---|
| | **t** | **df** | **p** | **95% conf (low)** | **95% conf (high)** | **cor** |
| branchLengthHuman | -6.8293 | 234 | 7.29E-11 | -0.51 | -0.30 | -0.41 |
| scaffoldN50 | 6.1701 | 234 | 2.98E-09 | 0.26 | 0.48 | 0.37 |
| contig N50 | 4.4414 | 234 | 1.38E-05 | 0.16 | 0.39 | 0.28 |
| **ANOVA** | | | | | | |
| **All samples with scaffold, contig N50, and branchLength to human vs annotated gene count** | | | | | | |
| Model: anova_test(HQGenes ~ scaffoldN50 + contigN50 + branchLengthHuman, data = cons) | | | | | | |
| Effect | DFn | DFd | F | p | ges | p<.05 |
| branchLengthHuman | 1 | 2.32E+02 | 39.096 | 1.92E-09 | 0.144 | * |
| scaffoldN50 | 1 | 2.32E+02 | 35.03 | 1.16E-08 | 0.131 | * |
| contigN50 | 1 | 2.32E+02 | 2.752 | 9.80E-02 | 0.012 | |

**Table S8.**

Olfactory receptor gene counts for 249 placental mammal species

**Table S9.**
Gene ontology analysis of genes whose protein-coding differences from the Eutherian mammal common ancestor are associated with the evolution of hibernation according to GLS forward genomics.

**A. Regions more conserved (pFDR <0.01) in deep hibernators than in strict homeotherms.**

| locus | Gene | Forward (f)/ Reverse (r) strand | Num trait loss species | # trait preserving species | Perfect match margin | GLS p-value | GLS adjusted p-value |
|---|---|---|---|---|---|---|---|
| chr1:68444529-68444685 | RPE65 | r | 154 | 22 | -0.06 | 1.04E-08 | 0.00045 |
| chr1:20654221-20654374 | DDOST | r | 153 | 22 | -0.06 | 1.11E-08 | 0.00045 |
| chr1:154552088-154552185 | UBE2Q1 | r | 148 | 22 | -0.09 | 2.75E-08 | 0.00088 |
| chr1:156477006-156477219 | MEF2D | r | 140 | 22 | -0.09 | 5.75E-08 | 0.00154 |
| chr1:12580521-12580677 | DHRS3 | r | 153 | 22 | -0.08 | 1.44E-07 | 0.00314 |
| chr1:77965066-77965238 | FUBP1 | r | 153 | 20 | -0.07 | 1.78E-07 | 0.00314 |
| chr1:68429768-68429941 | RPE65 | r | 153 | 22 | -0.12 | 1.98E-07 | 0.00314 |
| chr1:31908411-31908972 | PTP4A2 | r | 146 | 19 | -0.13 | 2.06E-07 | 0.00314 |
| chr10:830862-830987 | LARP4B | r | 153 | 22 | -0.05 | 2.34E-07 | 0.00314 |
| chr1:145995139-145995305 | TXNIP | r | 150 | 22 | -0.10 | 2.54E-07 | 0.00314 |
| chr1:20639884-20639997 | PINK1 | f | 151 | 22 | -0.12 | 2.80E-07 | 0.00321 |
| chr1:145994538-145994802 | TXNIP | r | 151 | 22 | -0.12 | 5.93E-07 | 0.00635 |
| chr1:12568339-12568426 | DHRS3 | r | 154 | 22 | -0.14 | 7.36E-07 | 0.00739 |
| chr1:12009590-12009731 | MFN2 | f | 152 | 22 | -0.08 | 1.01E-06 | 0.00954 |

**B. Significant (pFDR<0.05) overlaps with 7481 GO Biological Process gene sets ([www.gsea-msigdb.org/gsea/msigdb/annotate.jsp](www.gsea-msigdb.org/gsea/msigdb/annotate.jsp)) (*264*)**

| Gene Set Name | # Genes in Gene Set (K) | # Genes in Overlap (k) | k/K | p-value | FDR q-value | PMID |
|---|---|---|---|---|---|---|
| GOBP POSITIVE REGULATION OF MITOPHAGY IN RESPONSE TO MITOCHONDRIAL DEPOLARIZATION | 9 | 2 | 0.22 | 2.43E-06 | 1.82E-02 | 18200046, 23985961 |
| GOBP REGULATION OF AUTOPHAGY OF MITOCHONDRION IN RESPONSE TO MITOCHONDRIAL DEPOLARIZATION | 14 | 2 | 0.14 | 6.15E-06 | 2.30E-02 | 22020285 |
| GOBP RESPONSE TO MITOCHONDRIAL DEPOLARISATION | 20 | 2 | 0.1 | 1.28E-05 | 3.20E-02 | |
| GOBP MITOPHAGY | 24 | 2 | 0.08 | 1.86E-05 | 3.48E-02 | 15798367 |

**Supplementary data files**

**Data S1.** (separate file)
Outputs from GREAT analyses of ultraconserved elements (zooUCEs) and 100kb bins showing significant constraint.

**Data S2.** (separate file)
Input data for analysis of constraint in 100kb bins across the human genome. chr: Chromosome window is located on; start: start position of window; end: end position of window; zoonomiaBases: number of bases within window with phyloP scores; zoonomiaConsBases: Number of bases within window with phyloP > 2.27 (5% FDR constraint positions); zoonomiaNspeciesLow: Number of bases within window with a low (≤24) number of species aligning; zooN, A, C, G, T: Number of each base within window; zooFrac: Fraction of bases in window under constraint (phyloP > 2.27); k24gt90.sum: Number of bases with k24>0.9 (K24 UCSC track, a measure of mappability); cds.distinct.sum: Number of bases in CDS; cCRE.sum: Number of bases in cCREs; dhs.sum: Number of bases overlapping DHS.

**Data S3.** (separate file)
Hibernation RERConverge results with p-values to indicate whether a gene's relative evolutionary rate is associated with hibernation. Results when including (sheet 1) and excluding (sheet 2) bats (Chiroptera) are provided. Genes "In DEGs" are genes that have differential gene expression during different parts of the hibernation cycle (*87*)
